# When self comes to a wandering mind: Brain representations and dynamics of self-generated concepts in spontaneous thought

**DOI:** 10.1101/2021.12.18.473276

**Authors:** Byeol Kim, Jessica R. Andrews-Hanna, Jihoon Han, Eunjin Lee, Choong-Wan Woo

## Abstract

Self-relevant concepts are major building blocks of spontaneous thought, and their dynamics in a natural stream of thought are likely to reveal one’s internal states important for mental health. Here we conducted an fMRI experiment (*n* = 62) to examine brain representations and dynamics of self-generated concepts in the context of spontaneous thought using a newly developed free association-based thought sampling task. The dynamics of conceptual associations were predictive of individual differences in general negative affectivity, replicating across multiple datasets (*n* = 196). Reflecting on self-generated concepts strongly engaged brain regions linked to autobiographical memory, conceptual processes, emotion, and autonomic regulation, including the medial prefrontal and medial temporal subcortical structures. Multivariate pattern-based predictive modeling revealed that the neural representations of valence became more person-specific as the level of perceived self-relevance increased. Overall, this study provides a hint of how self-generated concepts in spontaneous thought construct inner affective states and idiosyncrasies.

## Introduction

> *“In the activity of association there is mirrored the whole psychical essence of the past and of the present, with all their experiences and desires. It thus becomes an index of all the psychical processes which we have but to decipher in order to understand the complete man*.*”*
>
> *by Eugen Bleuler*^1^

Our thoughts continuously come and go, transitioning from one to another topic even in the absence of overt task demands. This constant change and continuous flow are the key features of spontaneous thought^2-4^. Previous studies have found that adults spend a considerable amount of time engaging in spontaneous, perceptually-decoupled thought, a phenomenon commonly referred to as mind-wandering^5-7^. Since William James featured “the stream of thought” as a major subject of psychology^8^, researchers have posited the contents and dynamics of spontaneous thought to be important factors that can explain personality traits and mental health^2,9-11^. For example, recurrent negative thoughts are considered a transdiagnostic phenomenon given their links to many mood and anxiety disorders^12-14^. Adopting a complex dynamic systems view, spontaneous thought could be considered a random walk on a semantic network, in which the nodes represent autobiographical and semantic concepts, and the edges are the associations among the nodes established through past experiences^15-19^. In this framework, recurrent thoughts can be regarded as sticky nodes or strong attractors of the network^20-25^.

Self-relevant concepts are well-known attractors of spontaneous thought. Previous studies that examined the contents of spontaneous thought have found that the spontaneous thought contents are by no means random^2,3,26^. Rather, thought content tends to be self-relevant in nature, encompassing personal concerns, past memories, personal goals and planning, thinking about close others, etc.^3,16,27-31^ Based on these observations, a number of studies have suggested that self-referential processes are key functions of spontaneous thought^11,28,29,32-36^, and these findings were consistent across cultures^34,37,38^. Moreover, self-related spontaneous thought is known to be important for long-term health and psychological well-being^3,6,38-41^.

The past decade has brought increased interest in the neuroscience of spontaneous and task-unrelated thought, revealing associations with the brain’s default mode network (DMN)^42^. Though the DMN was initially featured as a signature of the brain’s “resting state”^43^, it is now more broadly appreciated for its role in multiple internally-guided cognitive and affective processes spanning spontaneous thought^44,45^, conceptual processing^46^, memory and future thinking^47,48^, mentalizing^49^, self-referential processes^50^, and autonomic and visceral modulation^51,52^. Collectively, these findings suggest that our brain is dynamically and continuously constructing our mental and bodily life. The quantitative assessment of brain representations and cognitive underpinnings of these dynamic processes would therefore help us better understand how and why the brain generates particular patterns of spontaneous thought, resulting in a healthy or unhealthy body and mind. In the current study, we examined the spontaneous thought dynamics and their brain representations using functional Magnetic Resonance Imaging (fMRI).

Despite the significance of dynamic, spontaneous thought in human psychology and psychopathology^53^, few quantitative tools and methods are currently available for neuroimaging studies. To overcome this challenge, we adapted a task we recently developed—called the Free Association Semantic Task (FAST)^54^—to use in conjunction with fMRI. The FAST integrates ideas from free word association^55,56^, experience sampling^57-61^, and naturalistic tasks^62-65^, and when adapted to a neuroimaging context, shows promise in revealing the dynamically unfolding signatures of spontaneous thought. The history of using free word association as a psychological test goes back to Francis Galton^66^, Wilhelm Wundt^67^, Emil Kraepelin^67^, and Carl Jung^68^, but modern psychology has largely ignored the method due to its questionable validity. However, the free association method has recently begun to receive attention again^54,56,69^, especially for its potential to be combined with computational methods, such as natural language processing and dynamic modeling to offer quantitative metrics into the emergence and unfolding of thought. Though early word association tasks recorded only one or two words in response to a seed word^66-68^, here we used a “chain” free association to better evaluate the dynamic characteristics of the stream of thought^56,70^. We focused on the affective and personal aspects of participants’ responses because emotionally charged and self-relevant thought topics are major ingredients of mind wandering and spontaneous thought^6,16,28^, and the free association method is known to be effective in revealing an individual’s emotional and autobiographical concepts^54,66,68^. Thus, we expect the FAST to provide a new way to probe the dynamic characteristics of affective and self-relevant thoughts, revealing personal inner affective states and idiosyncrasies that are important for human behaviors and mental health.

More specifically, this study aims to answer the following two research questions (**Fig. 1a**). First, are dynamic characteristics of spontaneous thought assessed with the FAST predictive of individual differences in emotional traits such as negative affectivity? Second, can we identify and decode the brain representations and dynamics of spontaneous thought? To answer these questions, we conducted an fMRI experiment (*n* = 62) in which participants completed the FAST involving three distinct phases (**Fig. 1b**). The first phase involved a “concept generation” phase, in which we asked participants to generate a concept as a word or phrase that came to their mind associated with the previous response every 2.5 seconds starting from a given seed word, leading to a self-generated associative concept chain. The responses were collected through an MR-compatible microphone, and participants generated a total of 40 consecutive concepts for each seed word. The second phase involved “concept reflection”, in which participants reflected on pairs of their self-generated concepts in sequence while undergoing fMRI scanning. This phase aimed to bring to mind the nature of the associative linkage between concepts generated by reflecting on the personal meaning of the concepts. The third part was the “post-scan survey.” After the fMRI scans, participants viewed their self-generated concepts once more and rated each concept using a multi-dimensional content scale^28^, which consisted of items evaluating emotional valence (how positive or negative is the concept?), self-relevance (how much is the concept relevant to yourself?), time (is the concept related to the past, present, or future?), vividness (does the concept involve vivid imagery?), and safety-threat (how much is the concept safe or threatening?). For more details of the task procedure, please see **Methods**.

**Figure 1.**
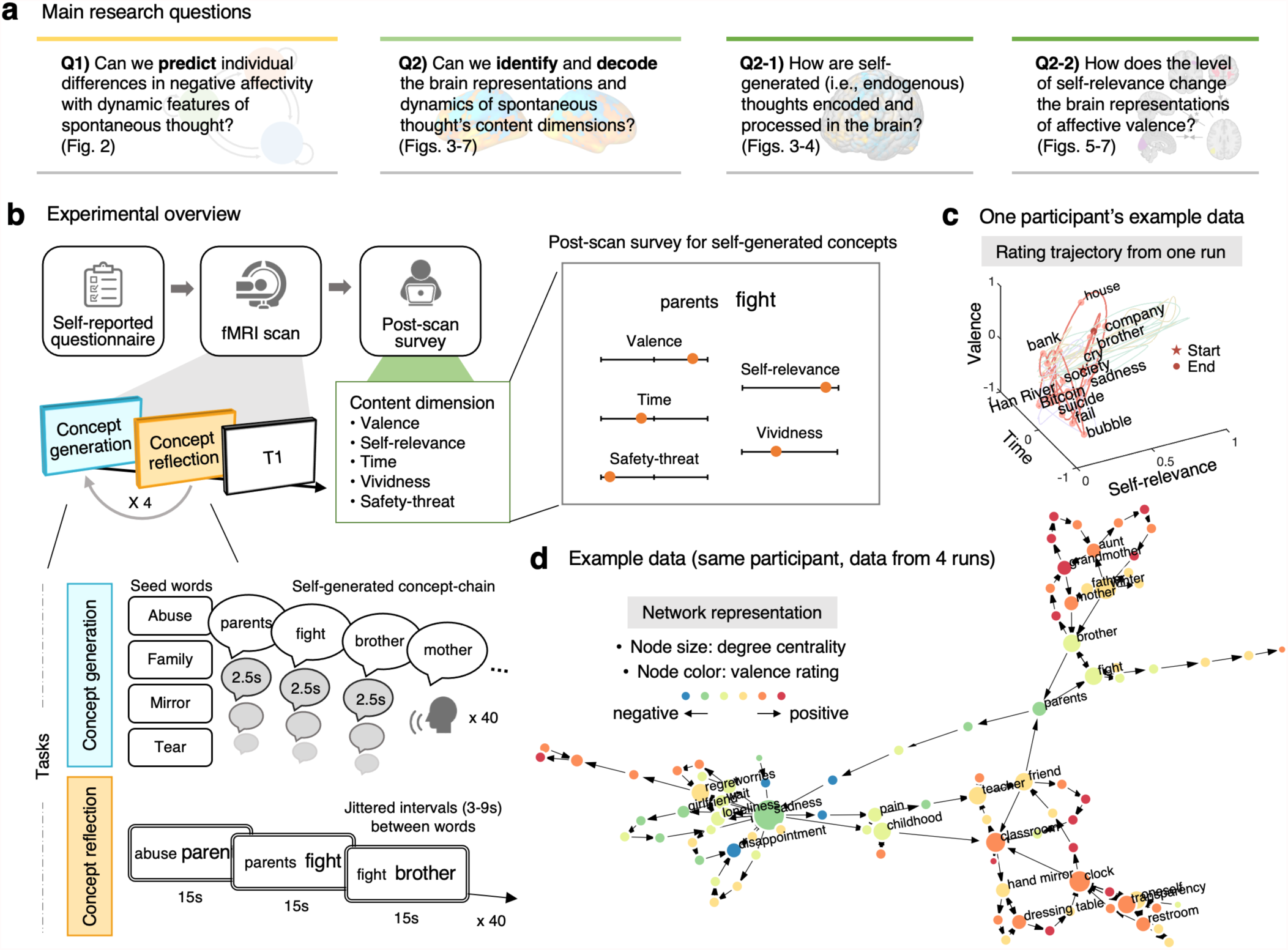
Research questions and experimental overview. **a**, Main research questions and their corresponding result figures. **b**, An experimental overview. Participants completed a battery of self-report questionnaires before the fMRI scans. During the fMRI scans, participants underwent the Free Association Semantic Task (FAST), which consisted of three main parts—concept generation, concept reflection, and post-scan survey. The fMRI experiment had four runs, each of which included the concept generation and concept reflection tasks. For the concept generation task, we asked participants to report a word or phrase that came to their mind in response to the previous concept every 2.5 seconds starting from a given seed word. The seed words for the first session included “family”, “tear”, “mirror”, and “abuse.” After participants generated 40 concept-chain responses, we showed the two consecutive responses and asked them to think about the target (i.e.., the second) concept’s personal meaning for 15 seconds. After the scan, we asked participants to complete a post-scan survey on the 160 self-generated concepts. We showed the two consecutive concepts again and asked participants to rate the target concept in terms of their valence, self-relevance, time, vividness, and safety-threat. **c**, One participant’s example data shown on the 3-dimensional space of valence, self-relevance, and time. The dots indicate self-generated responses, and the pink line indicates data from one run. A red star indicates the start of the run, and the red dot indicates the ending point. **d**, A network representation of one participant’s data. The dots represent the self-generated responses, and the arrows show the direction of the concept generation. The dot colors represent the averaged valence scores of each concept, and the dot size indicates the degree centrality (i.e., how many edges are connected to the node).

Markov chain analyses combined with machine learning on the post-scan survey data revealed that a prediction model based on the temporal dynamics of FAST responses was highly predictive of individual differences in negative affectivity (*r* = 0.486), and further generalized to three independent test datasets (total *n* = 196, *r* = 0.492-0.628). While participants reflected on the personal meaning of self-generated concepts, medial prefrontal and medial temporal brain structures were strongly activated. To identify the brain representations of some phenomenological characteristics of the self-generated concepts, we trained fMRI pattern-based predictive models of the content dimensions using the fMRI data from the concept reflection phase. Among the multiple content dimensions, we achieved a significant prediction only for the self-relevance dimension (*r* = 0.305), but for the valence dimension, we were able to develop well-performing predictive models (*r* = 0.307) when we used only the trials with low self-relevance. We further found that the brain representations of valence became more idiosyncratic (i.e., person-specific) when the self-generated concepts were rated high in self-relevance.

Overall, the current study developed a new experimental method that allows us to examine the dynamic characteristics of spontaneous thought and their brain representations, paving the way for quantitative modeling of spontaneous thought dynamics. Additionally, our findings provide a deeper understanding of where in the brain self-generated, endogenous thoughts are represented and how self-relevance modulates the brain’s affective representations.

## Results

### Dynamics of spontaneous thought probed with the Free Association Semantic Task

**Figs. 1c-d** shows data from a representative participant to illustrate how FAST responses can reveal the characteristic topics and dynamic features of an individual’s spontaneous thought. As shown in **Fig. 1c**, FAST responses can be viewed in the context of a high-dimensional state space of some phenomenological characteristics. This participant’s concept association initially flowed from a given seed word, ‘tear,’ to negatively valenced concepts (‘cry,’ ‘sadness,’ and ‘suicide’), then moved to societal and less self-relevant thought topics (‘society,’ ‘Bitcoin,’ ‘bubble,’ ‘fail,’ and ‘bank’) and finally transitioned to personal topics (‘house,’ ‘company,’ and ‘brother’). This example highlights that the FAST can reveal topics of spontaneous thought that ranges from personal narratives to societal events and issues (e.g., the data were collected early 2018 when the Bitcoin crash occurred).

FAST responses can also be viewed as a directed graph in which the nodes are the response words, and the directed edges are the connections from the previous concepts to the next associative concepts (**Fig. 1d**). Key hubs of this participant’s graph, including ‘sadness,’ ‘brother,’ and ‘classroom,’ can be seen as an attractor that is densely connected to other related concepts. FAST responses tended to come back to these nodes as if these had strong gravity in this personal semantic state space.

### Free association dynamics predictive of individual differences in negative affectivity

To answer our first research question (Q1 in **Fig. 1a**, i.e., whether the affective dynamics of spontaneous thought assessed with the FAST are predictive of individual differences in negative affectivity), we used the Markov chain analysis to create input features for machine learning to build a predictive model of general negative affectivity based on the transitional dynamics on the content dimensions. Prior to the analysis, we ensured that the post-scan survey ratings reflected participants’ in-scanner experience by showing that 1) the in-scanner heart-rates were significantly modulated by the levels of post-scan valence ratings, and 2) the valence ratings were consistent with the emotion ratings intermittently obtained during the fMRI scans (see **Supplementary Figs. 1** and **2** for detail).

For the Markov chain analysis, we first defined discrete states by dividing each content dimension into two or three discrete states, as shown in **Fig. 2a**. We divided valence, time, and safety-threat, which ranged from -1 to 1, into three discrete states (−1 to -0.33, -0.33 to 0.33, and 0.33 to 1; for valence, the three discrete states were negative/neutral/positive, for time, past/present/future, for safety-threat, threatening/neutral/safe). For the self-relevance and vividness dimensions that ranged from 0 to 1, we divided them into two discrete states (0 to 0.5 and 0.5 to 1, which corresponded to low and high for both dimensions, respectively). We then calculated the transition probability, which was defined as the probability of making transitions from one to another discrete state on each dimension. We also calculated the steady state probability, which was defined as the probability of converging to one state when the transition processes were sufficiently repeated^71^. In addition to these dynamic features from the Markov chain analysis, we also used each affective dimension’s mean and variance as predictor variables for the subsequent predictive modeling. Many of these dynamic features were relatively stable over a 7-week interval (for their test-retest reliability, see **Supplementary Table 1**).

**Figure 2.**
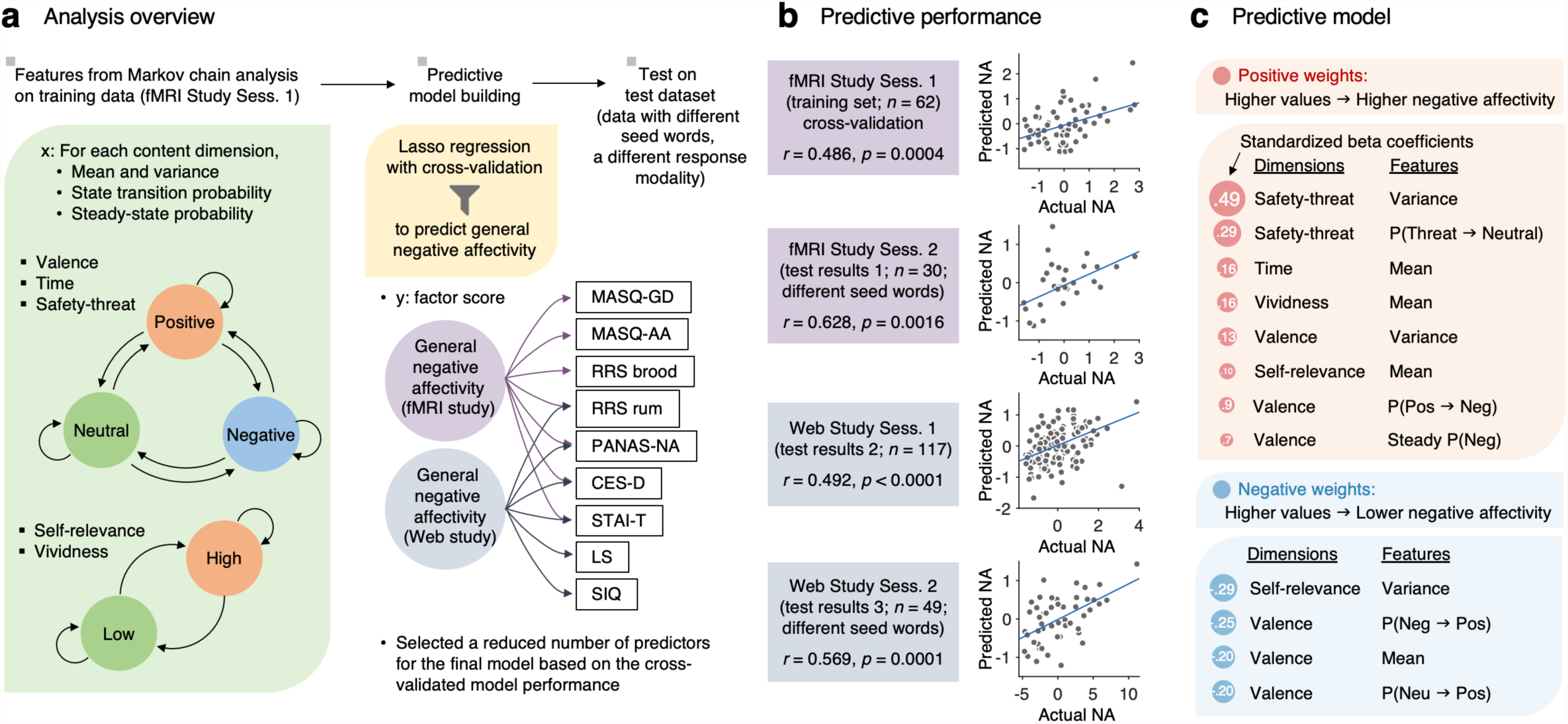
Markov chain-based predictive modeling of negative affectivity. **a**, Analysis overview. The input features for the predictive modeling included the state transition and steady-state probabilities estimated with the Markov chain analysis and the mean and variance of the content dimension ratings. For the Markov chain analysis, we defined three discrete states for valence, time, and safety-threat, which ranged from -1 to 1, and two discrete states for self-relevance and vividness, which ranged from 0 to 1. With these features, we developed predictive models of general negative affectivity, which we modeled with factor analyses. The fMRI and web studies included the overlapping set of questionnaires, but they were not the same. Thus, we built two separate factor models—one for the fMRI study, and the other for the web study. Though the actual factor models also included self-report questionnaires for a factor of general positive affectivity, here we show the questionnaires only for the general negative affectivity factor for brevity. For the details of the factor analysis results, please refer to **Supplementary Table 2**. We used a Lasso regression as a fitting algorithm. **b**, Model performance. From top to bottom, the plots show 1) the leave-one-participant-out cross-validated prediction results within the training dataset (*n* = 62, first session of the fMRI study), and three independent test results on 2) the second session re-test data of the fMRI study with different seed words (*n* = 30), 3) the first session data of the FAST-web study (*n* = 117), and 4) the second session re-test data of the FAST-web study with different seed words (*n* = 49). The actual versus predicted negative affectivity factor scores are shown in the plots. Each dot represents each participant. We evaluated the model performance with robust correlation between the actual and predicted levels of general negative affectivity. NA, negative affectivity. **c**, To interpret the final model, we examined the standardized beta coefficients of the input features of the model. From the lasso regression, a total of 12 features were selected. The features in red indicate positive weights, whereas the features in blue indicate negative weights. MASQ-GD, Mood and Anxiety Symptom Questionnaire-General Distress; MASQ-AA, MASQ-Anxiety arousal; RRS brood, Rumination Response Scale-Brooding; RRS rum, RRS-Depressive rumination; PANAS-NA, Positive and Negative Affect Schedule-Negative Affect; CES-D, Center for Epidemiologic Studies Depression; STAI-T, State-trait anxiety inventory-Trait version; LS, Loneliness Scale; SIQ, Suicidal Ideation Questionnaire.

For the predictive modeling, we used Lasso regression^72^ to predict individual differences in general negative affectivity with leave-one-subject-out cross-validation. The number of predictor variables for the final model was determined based on the cross-validation performance with the training data (*n* = 62; one participant was excluded due to excessively few responses). We then tested the final model on three testing datasets to evaluate two different generalizability types—seed words and response modality. First, we tested the model on re-test data with an average 7-week interval, in which a different set of seed words were used on a subset of participants from the training dataset (*n* = 30). Second, we tested the model on an independent test data (*n* = 117) collected from a new set of participants using a web-based FAST experiment, which collected the response through typing instead of speaking and with a longer time limit for a response (for more detail of the web-based FAST, please see **Methods**). Third, we tested the model on re-test web-based FAST data with an average 7-week interval, in which, again, a different set of seed words were used on a subset of participants from the web-based FAST dataset (*n* = 49). As the outcome variable, we used factor scores from a factor analysis of sub-scales from self-report questionnaires measuring multiple aspects of general negative affectivity.

As shown in **Fig. 2b**, the final predictive model showed significant prediction performance across four datasets; for the training dataset (*n* = 62) with leave-one-subject-out cross-validation, the prediction-outcome correlation between the actual and predictive values was *r* = 0.486, *p* = 0.0004, two-tailed, one-sample *t*-test, for the re-test data (*n* = 30) with different seed words, *r* = 0.628, *p* = 0.0016, for the web-based FAST experiment (*n* = 117), *r* = 0.492, *p* < 0.0001, and for the web-based FAST re-test (*n* = 49) with different seed words, *r* = 0.569, *p* = 0.0001. Given that these three independent test datasets had slightly different experimental parameters, such as seed words and response modality, these results demonstrated the robustness of the task and the predictive model.

We then examined the standardized beta coefficients of the 12 behavioral dynamic features to determine which predictors contributed significantly to the final model of general negative affectivity (**Fig. 2c**). The beta coefficients indicated that the participants who showed 1) a higher variance of the safety-threat and valence scores, 2) a higher transition probability from threatening to neutral states, 3) a higher mean score on the time, vividness, and self-relevance dimensions (i.e., relatively more future-oriented, more vivid, and more self-relevant), 4) a higher transition probability from positive to negative states, and 5) a higher steady-state probability for the negative state were likely to report a higher level of general negative affectivity. Conversely, participants who showed 1) a higher variance on the self-relevance score, 2) a higher transition probability from the negative or neutral to positive states, and 3) a higher mean score on the valence dimension (i.e., more positive) tended to report a lower level of general negative affectivity.

We also trained an additional predictive model with a subset of the content dimension— valence, self-relevance, and time, and thus we called this reduced model a VST model—to see whether these three dimensions were enough to predict the level of general negative affectivity. We chose these three dimensions because valence and self-relevance were highly correlated with safety-threat and vividness, respectively (**Supplementary Fig. 3a**). The VST model also showed significant predictions across four datasets with *r* = 0.409 to 0.677, *p* = 0.0042 to *p <* 0.0001, and seven out of eight selected features overlapped with the original full model features (**Supplementary Fig. 4**).

**Figure 3.**
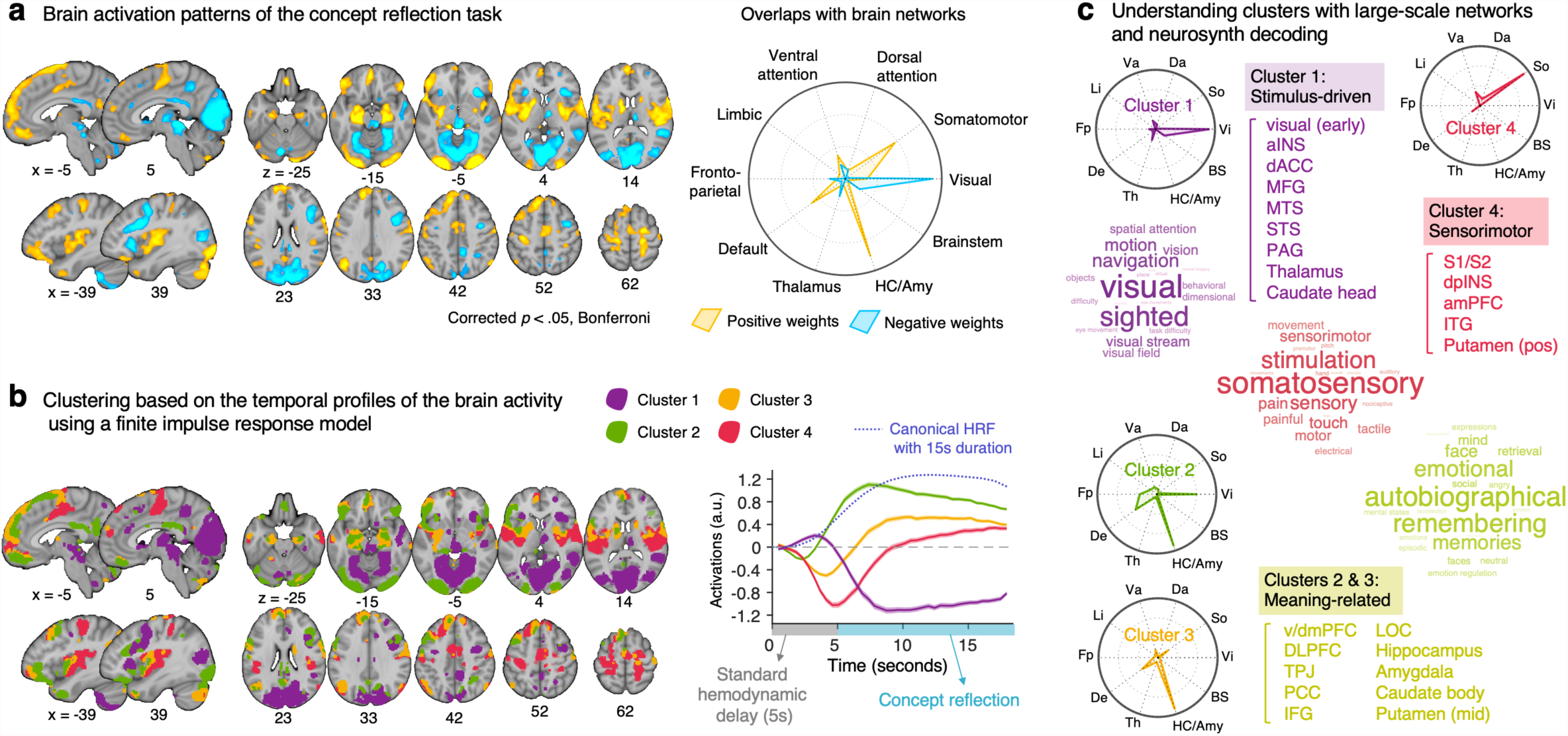
Brain activation patterns during the concept reflection task. **a**, The basic contrast map for concept reflection versus baseline thresholded at *p* < 0.05 with Bonferroni correction. The radial plot shows the relative proportions of overlapping voxels between the thresholded map and large-scale networks (or regions) given the total number of voxels within each network (or each region). For the definitions of large-scale networks and regions, see **Supplementary Fig. 5. b**, To better understand the temporal patterns of brain activity during concept reflection, we conducted a clustering analysis based on a finite impulse response model. We identified four clusters, and the plot on the right shows the time-course of averaged activity across voxels of each cluster. Shading represents the standard error of the mean (s.e.m.). On the x-axis, zero represents the onset of the trial, and the gray area represents the standard hemodynamic delay, which peaks at around 5 seconds. The blue dotted line shows the canonical hemodynamic response function (HRF) for an event with a 15-second duration. **c**, To interpret the functional meaning of the clusters, we examined the clusters with the large-scale functional networks and the term-based decoding analysis based on an automated large-scale meta-analysis, neurosynth^73^. We combined the clusters 2 and 3 because they showed similar profiles in terms of their time-course and anatomical locations. Word clouds show the term-based decoding results, and the font size indicates the relative relevance with the term. For the radial plot, Va, ventral attention; Da, dorsal attention, So, somatomotor; Vi, visual; BS, brainstem; HC/Amy, hippocampus/amygdala; Th, thalamus; De, default; Fp, frontoparietal; Li, limbic. For the brain regions, v/dmPFC, ventral/dorsal medial prefrontal cortex; DLPFC, dorsolateral prefrontal cortex; TPJ, temporal parietal junction; PCC, posterior cingulate cortex; IFG, inferior frontal gyrus; LOC, lateral occipital cortex; S1/S2, primary and secondary somatosensory cortex; dpINS, dorsal posterior insula; amPFC, anterior medial prefrontal cortex; ITG, inferior temporal gyrus; aINS, anterior insula; dACC, dorsal anterior cingulate cortex; MFG, middle frontal gyrus; MTS, middle temporal sulcus; STS, superior temporal sulcus; PAG, periaqueductal gray. For the locations of each region, please see **Supplementary Fig. 6**.

**Figure 4.**
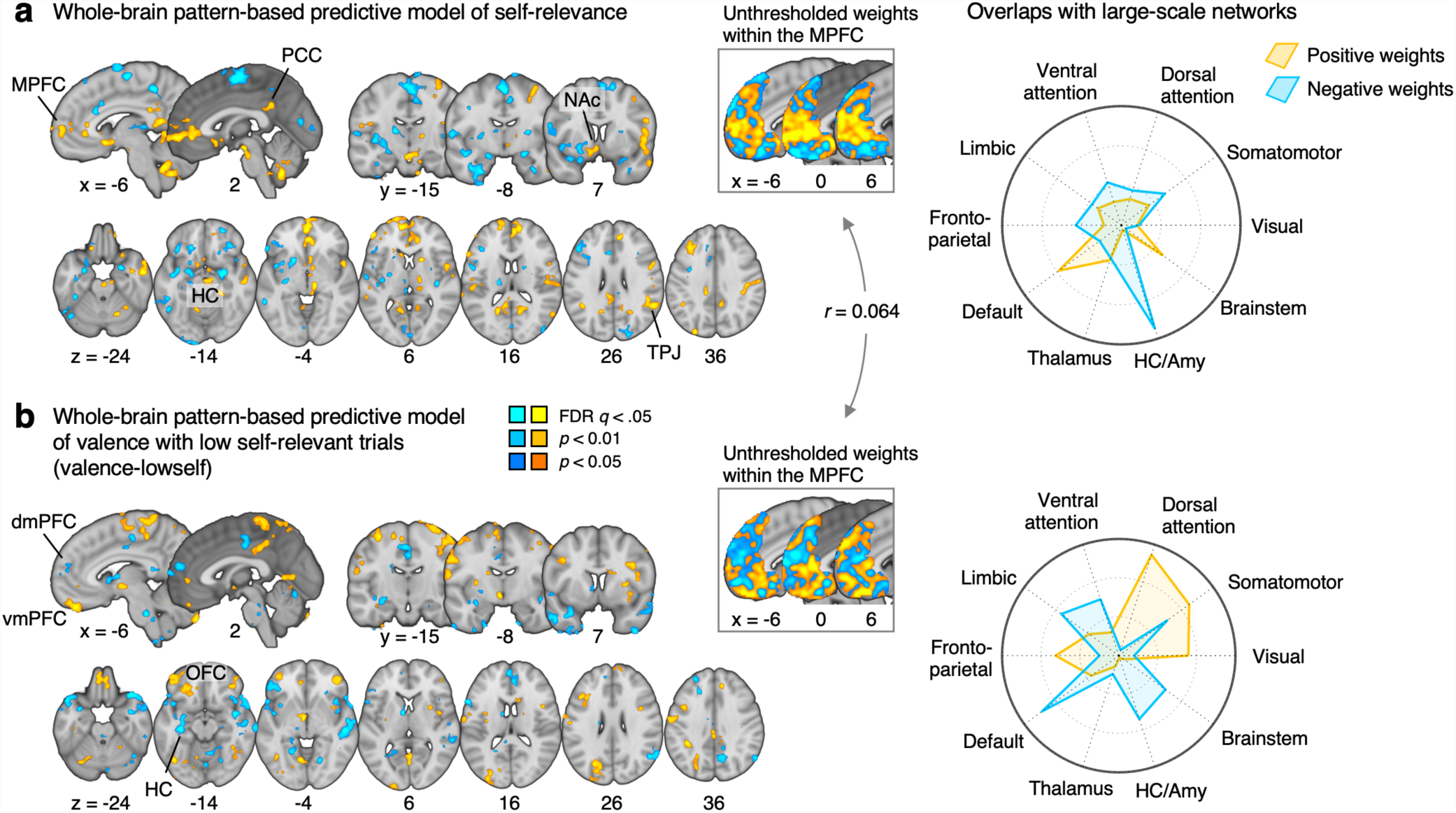
Multivariate pattern-based predictive models of self-relevance and valence. **a**, Self-relevance model. The map shows the voxels that reliably contributed to the prediction of self-relevance scores based on bootstrap tests (thresholded at FDR *q* < 0.05, two-tailed). Thresholding was performed for the display purpose; all weights were used in the prediction. We also pruned the map using two additional more liberal thresholds, uncorrected *p* < .01 and *p* < .05, two-tailed, to show the extent of activation clusters. The radial plot shows the relative proportions of overlapping voxels between the thresholded map and large-scale networks given the total number of voxels within each network. **b**, Valence model for the trials with low self-relevance scores (valence-lowself). Insets show the unthresholded predictive weights within the medial prefrontal cortex (MPFC) of the two models. The correlation value (*r* = 0.064) indicates the pattern similarity of the MPFC weights between the two models. PCC, posterior cingulate cortex; TPJ, temporal parietal junction, NAc, nucleus accumbens; vmPFC, ventromedial prefrontal cortex; dmPFC, dorsomedial prefrontal cortex; OFC, orbitofrontal cortex; HC, hippocampus.

### Brain activation patterns during the concept reflection phase

To answer our second research question (Q2-1 in **Fig. 1a**; identifying the brain representations of the content dimensions of spontaneous thought), we first examined brain activation patterns while participants reflected on the self-generated concepts in the context of their conceptual associations. As shown in **Fig. 3a**, brain regions spanning the hippocampus, amygdala, and parts of the somatomotor network and default mode network engaged to a greater degree during conceptual self-reflection (warm color) compared to fixation baseline. In contrast, the visual network was engaged to a greater degree during baseline than during conceptual self-reflection, and thus appeared “deactivated” during reflection (see **Supplementary Fig. 5** for the large-scale network definitions used for identification purposes).

**Figure 5.**
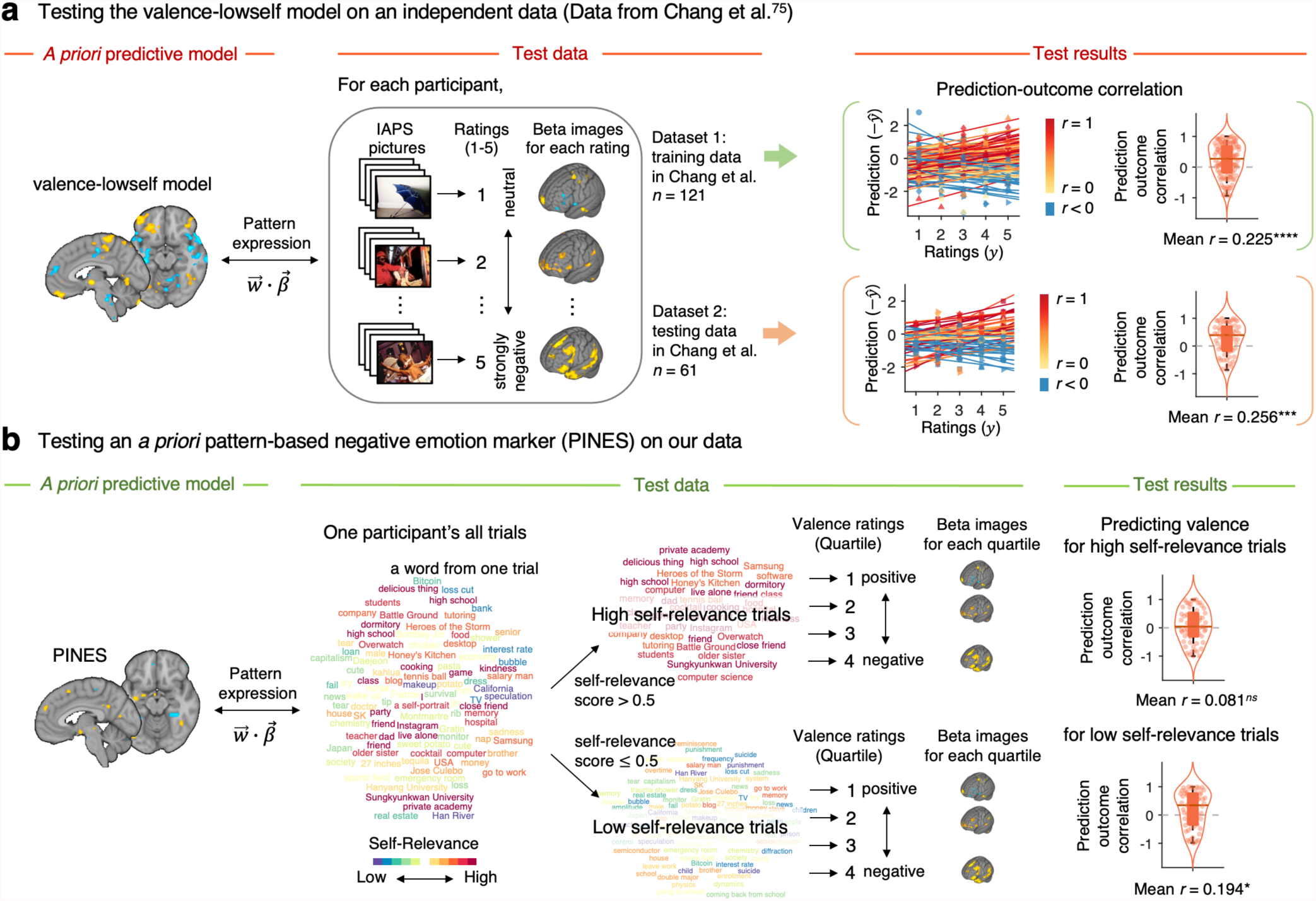
Cross-testing of two *a priori* models of valence on two independent datasets. **a**, To further validate our valence-lowself model, we tested the model on an independent study dataset, in which negative emotions were induced using the International Affective Picture System (IAPS) pictures^78^. To this end, we applied the valence-lowself model on the beta images from the independent dataset corresponding to five-points negative emotion ratings ranging from 1 (neutral) to 5 (strongly negative). We obtained predicted ratings by calculating a dot product of each vectorized test image data with the model weights. The plot on the right shows the actual versus predicted ratings, and for convenience, we added the negative sign to the predicted ratings to make the expected prediction into positive correlations. Each colored line in the plot represents individual participant’s data (red, higher *r*; yellow, lower *r*; blue, *r* < 0). The violin and box plots display the distributions of within-participant prediction-outcome correlations.^***^*p* < 0.001, ^****^*p* < 0.0001, bootstrap tests, two-tailed. **b**, To test whether the brain representation of valence was modulated by the level of self-relevance, we tested an *a priori* neuroimaging emotion marker, Picture-Induced Negative Emotion Signature (PINES)^78^ on our data, separately for trials with high self-relevance scores (> 0.5) versus low self-relevance scores (≤ 0.5). The model performance was based on the quartile data based on the valence scores, and the violin and box plots on the right show the distributions of within-participant prediction-outcome correlations. ^*ns*^not significant, ^*^*p* < 0.05, bootstrap tests, two-tailed.

Further investigation into the temporal shape of these hemodynamic response patterns using a finite impulse response (FIR) model revealed that the visual cortex “deactivation” was driven by a transient increase in activity around 3 seconds after stimulus onset, followed by a large decrease afterwards (**Figs. 3b**). K-means clustering on the FIR signal across the brain showed that the visual cortex, some brain regions within the ventral attention network, and thalamus formed a cluster. This cluster (Cluster 1) showed a transient activity right after the stimulus onset, likely reflecting perceptually-guided and attentional orienting processes. Two additional clusters (Clusters 2-3) emerging from the clustering analysis mainly consisted of default mode and limbic network, lateral prefrontal cortex, and hippocampus and amygdala regions. Both clusters showed a delayed peak of brain activity around 7-10 seconds after the stimulus onset. Another cluster (Cluster 4) that had a large overlap with the somatomotor network showed a negative peak around 5 seconds after the stimulus onset. Given that the concept reflection did not involve any actual sensorimotor experience, this deactivation seemed reasonable. However, as shown in **Fig. 3b**, the brain activation level within this cluster showed a slow recovery and turned into positive activation towards the end of the trial.

To functionally characterize the four activity clusters, we examined our clustering results with the term-based decoding analysis based on 1) an automated large-scale fMRI meta-analysis (neurosynth.org^73^), 2) the principal gradient of cortical hierarchy^74^, and 3) basal ganglia parcellations^75^. As shown in **Fig. 3c**, the first clusters showed the largest correlations with the meta-analytic maps with functional terms including “visual,” “sighted,” and “navigation.” The second and third cluster showed the largest correlations with the terms “autobiographical,” “remembering,” and “emotional.” The fourth cluster were correlated with the terms “somatosensory,” “stimulation,” and “sensory.” Based on the decoding results, we combined and named the second and third clusters as “meaning-related” because many of the brain regions within these clusters have been shown to be involved in semantic processing and autobiographical memory. In addition, we named the first cluster “stimulus-driven” and the fourth cluster as “sensorimotor,” respectively (see **Supplementary Fig. 6** for the details of the regions included in each cluster).

**Figure 6.**
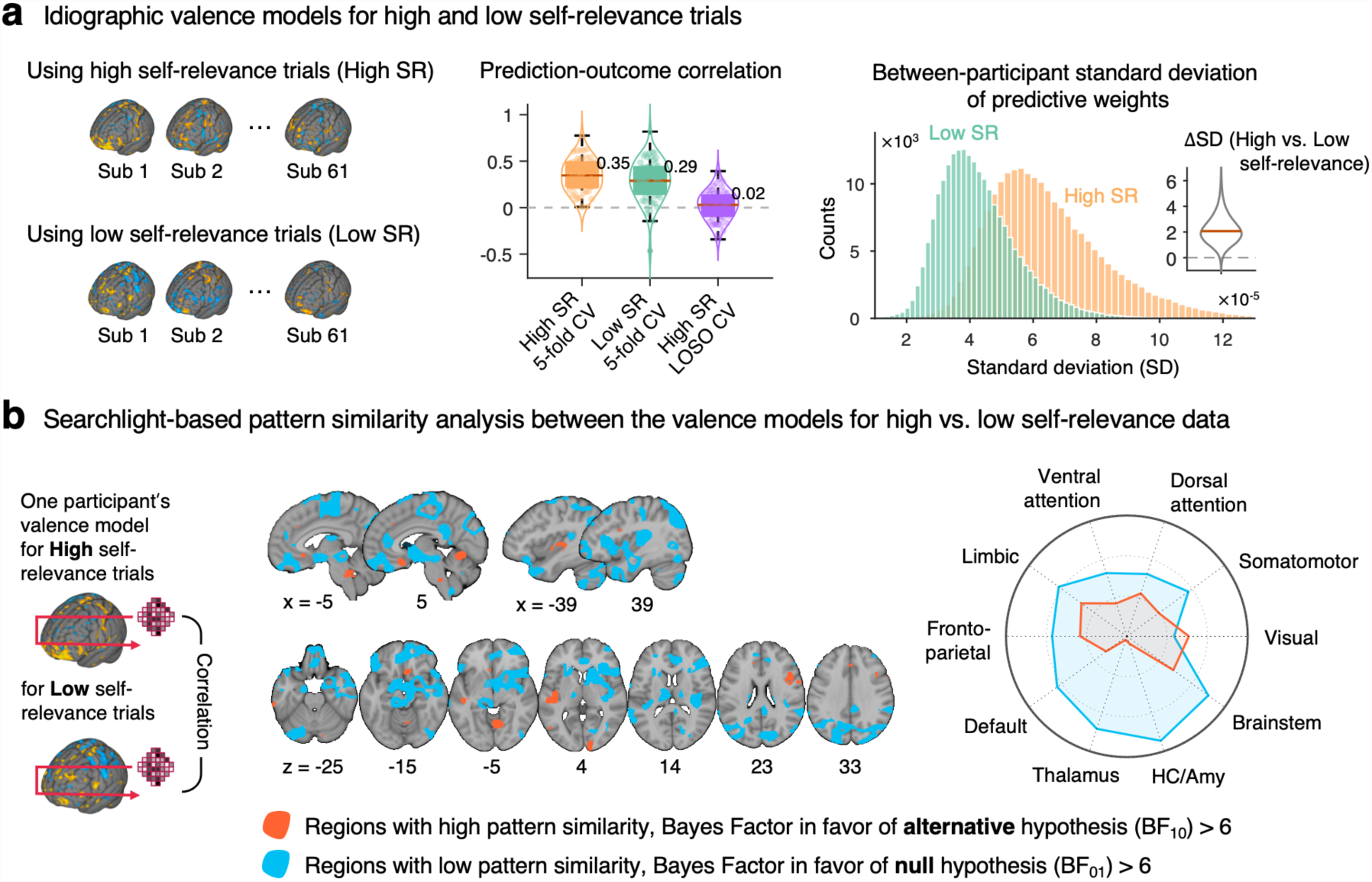
Idiographic predictive modeling of valence. **a**, We used an idiographic predictive modeling approach to quantify the between-participant variability of predictive weights for high versus low self-relevance data. We trained two valence models per person—the first model used data from the trials with the high self-relevance scores (> 0.5; High SR), and the other model used data from the trials with low self-relevance scores (≤ 0.5; Low SR). In addition, we trained a group-level valence model by concatenating all participants’ high self-relevance data with leave-one-subject-out cross-validation (LOSO-CV) for comparison. The violin and box plots in the middle show the prediction performance of two idiographic models with 5-fold CV and the group-level model with LOSO-CV. Each dot represents each participant. The histograms on the right show the distributions of between-participant standard deviation of predictive weights, and the inset violin plot shows the differences in the standard deviation between two valence models across voxels. **b**, We examined which brain regions displayed similar or distinct spatial patterns of predictive weights between two valence model with a searchlight-based pattern similarity analysis. For this, we first created a searchlight with a radius of 5 voxels and scanned it throughout the whole brain with a step size of 4 voxels. We then calculated correlation coefficients between two models’ predictive weights within each searchlight. The red regions showed Bayes factor values in favor of having distinct patterns between two models (BF_10_ > 6), whereas the blue regions were the opposite, i.e., in favor of having similar patterns (BF_01_ > 6). The radial plot shows the relative proportions of overlapping voxels between the thresholded map and large-scale networks given the total number of voxels within each network.

Furthermore, as shown in **Supplementary Fig. 7**, our region clustering and their naming were largely consistent with the principal gradient in the cortex and the meta-analysis findings in the basal ganglia. For example, our clusters named “meaning-related” largely overlapped with the trans-modal end in the principal gradient of cortical hierarchy (**Supplementary Fig. 7a**) and the parts of the basal ganglia related to social, language, and executive functions (**Supplementary Fig. 7b**). The “stimulus-driven” and “sensorimotor” clusters overlapped with the unimodal end of the cortical principal gradient and the basal ganglia parcellations for stimulus value and sensorimotor processes, respectively. These findings support that our clustering analysis resulted in neurobiologically meaningful clusters, providing a basis for further analyses of our data and functional interpretations of our findings.

### Multivariate pattern-based predictive models of self-relevance and valence

To further investigate our second research question (“Can we identify and decode the brain representations of affective qualities of spontaneous thought?” in **Fig. 1a**), we developed multivariate pattern-based predictive models for the content dimension ratings. To prepare training data, we grouped trials into quartiles representing four levels of each content dimension scale and then averaged the brain and rating data, resulting in four brain maps and four rating scores per person for each dimension. After concatenating all participants’ data (*n* = 61), we trained principal components regression (PCR) models for each content dimension and estimated model performance using two types of cross-validation methods--leave-one-subject-out cross-validation (LOSO-CV) and random-split cross-validation (RS-CV)^76,77^. The cross-validated prediction performance was significant for self-relevance (correlation between actual and predicted ratings: with LOSO-CV, mean *r* = 0.304, *z* = 4.400, *p <* 0.0001, two-tailed, bootstrap tests, mse = 0.155; with RS-CV, *r* = 0.276, mse = 0.156; **Fig. 4a**), while other dimensions showed relatively poor prediction performance, with LOSO-CV, mean *r* = 0.185, mse = 0.399 for valence; mean *r* = 0.166, mse = 0.319 for safety-threat; mean *r* = -0.064, mse = 0.228 for time; *r* = -0.015, mse = 0.182 for vividness. With RS-CV, mean *r* = 0.152, mse = 0.427 for valence; mean *r* = 0.147, mse = 0.323 for safety-threat; mean *r* = -0.012, mse = 0.223 for time; *r* = -0.002, mse = 0.183 for vividness.

Among the dimensions that showed poor prediction performance, the valence result was unexpected because previous studies have showed reasonable performance in predicting positive vs. negative emotional valence. For example, Chang et al.^78^ reported that a whole-brain pattern-based marker could predict negative emotion ratings induced by pictures with high prediction performance. Other studies also reported that regional brain activity patterns could classify the positive versus negative valence with significant classification accuracy^79,80^. However, unlike the previous studies, which used exogenous stimuli to evoke emotions, such as pictures^78^, movies^79^, or tastants^80^, the current study used self-generated, endogenous stimuli, which could have a potential impact on the semantic representations of emotional valence in the brain. Thus, we hypothesized that if we trained a predictive model only with the data from trials with low self-relevance scores, we might be able to achieve a significant prediction performance similar to the previous studies. To this end, we separately trained two models of valence, one for the low self-relevance trials (self-relevance scores ≤ 0.5), and the other for the high self-relevance trials (self-relevance scores > 0.5). Other analysis procedures were the same as previous.

As hypothesized, we found that the valence model trained only on the low self-relevance trials (named the “valence-lowself” model; **Fig. 4b**) showed a better and significant prediction performance, mean *r* = 0.307, *z* = 3.808, *p* < 0.0001, bootstrap test, mse = 0.362 with LOSO-CV, and *r* = 0.303, mse = 0.364 with RS-CV, than the valence model trained on the data with high self-relevance, mean *r* = 0.031, *z* = 0.403, *p* = 0.6872, mse = 0.448 with LOSO-CV, and *r* = 0.060, mse = 0.426 with RS-CV. To further validate this valence-lowself model, we tested the model on an independent study dataset from Chang et al.^78^. We chose this study dataset because it used exogenous emotional stimuli to induce emotions (i.e., the International Affective Picture System pictures) and was publicly available from NeuroVault (https://identifiers.org/neurovault.collection:503). As shown in **Fig. 5a**, when we applied the valence-lowself model on the beta images corresponding to five-points negative emotion ratings ranging from 1 (neutral) to 5 (strongly negative), our model showed significant predictions across two independent datasets. The first dataset was the training data in the original study^78^ (*n* = 121, mean *r* = 0.225, *p* < 0.00001, two-tailed, bootstrap tests), and the second dataset was the testing data in the original study (*n* = 61, *r* = 0.256, *p* = 0.0002). These results provided evidence for the generalizability of our valence-lowself model to emotions evoked with exogenous visual stimuli.

To further understand the neurobiological meaning and validity of the predictive models, we visualized the thresholded predictive maps of self-relevance (**Fig. 4a**) and valence-lowself (**Fig. 4b**) based on bootstrap tests with 10,000 iterations and the false discovery rate (FDR) *q* < 0.05, identifying brain voxels that made reliable contributions to the prediction. For the self-relevance model, multiple brain regions within the default mode and limbic networks appeared to be important, including the medial prefrontal cortex (MPFC), PCC, TPJ, TP, hippocampus, and nucleus accumbens (NAc), consistent with previous literature^31,49,81-85^. Similarly, the valence-lowself model also identified important predictors within the default mode and limbic networks, such as the dorsal and ventral medial prefrontal cortex (dm/vmPFC), orbitofrontal cortex (OFC), and hippocampus. However, the predictive weight patterns within these regions were quite different between the self-relevance and valence-lowself models. For example, as shown in the insets of **Fig. 4**, which presented the unthresholded weights of the self-relevance and valence-lowself models within the MPFC, the self-relevance model showed a negative → positive → negative gradient from dorsal to ventral parts of the MPFC. In contrast, the valence-lowself model showed a negative → positive gradient from dorsal to ventral MPFC. The pattern similarity of the unthresholded predictive weights within the MPFC between the two models was low, *r* = 0.064. In addition to the default mode and limbic network regions, many voxels within the somatomotor and ventral and dorsal attention networks were among the important features of the models, suggesting that the information about the levels of self-relevance and valence involves many brain regions distributed across multiple brain systems^78,86-90^.

Note that we did not further examine the predictive models of the other three dimensions, i.e., vividness, safety-threat, and time, given that the principal component analysis results suggested that there were three main principal components (PCs) in the content dimensions. As shown in **Supplementary Fig. 3a**, the valence and safety-threat dimensions were highly correlated, and the self-relevance and vividness were also highly correlated. Therefore, by modeling valence and self-relevance, we should be able to cover the first two PCs. Regarding the time dimension, its predictive model did not perform well, and thus we did not further examine the model here. In **Supplementary Fig. 3b**, we presented the univariate general linear model results with the three PCs.

### Idiosyncratic brain representations of emotional valence for high self-relevance trials

One of the intriguing observations in the previous section was the poor prediction performance of the valence model when it was trained on the high self-relevance trials. We hypothesized that valence information for the high self-relevance trials would be represented with more idiosyncratic brain activity patterns than the low self-relevance trials. To test the hypothesis, we first tested whether an *a priori* multivariate pattern-based emotion marker provided a similar pattern of results. We used the Picture-Induced Negative Emotion Signature (PINES)^78^, which has shown its sensitivity and specificity in predicting the level of negative emotions across multiple studies^78,91^. As presented in **Fig. 5b**, the results were consistent with our findings in the previous section—the PINES was able to predict the valence ratings only for the data from the low self-relevance trials, mean *r* = 0.194, *p* = 0.0171, two-tailed, bootstrap tests, but not for the high self-relevance trials, *r* = 0.081, *p* = 0.2813. These results suggest that the data from low self-relevance trials had some shared pattern information for valence common across participants, which could be captured by the PINES, whereas the valence information from high self-relevance trials cannot be decoded with the population-level emotion marker.

We then used an idiographic predictive modeling approach to quantify the between-subject variability of predictive weights for high versus low self-relevance data. We first split the trials into high and low self-relevance groups (using 0.5 as a cutoff score) and trained two valence models per person—one for high self-relevance trials and the other for low self-relevance trials using PCR with 5-fold cross-validation. Though the prediction performances were not different between the high versus low self-relevance predictive models (*t*_60_ = 1.886, *p* = 0.0641, two-tailed, paired *t*-test; the middle panel of **Fig. 6a**), the standard deviations of the predictive weights across participants were significantly higher in the high self-relevance models than the low self-relevance models (*t*_211362_ = 867.59, *p* < 0.0001; the right panel of **Fig. 6a**). Moreover, a group-level model trained on the high self-relevance data from all participants showed significantly worse prediction performance than the idiographic model built individually with the same data (*t*_60_ = 10.005, *p* < 0.0001, two-tailed, paired *t*-test; the middle panel of **Fig. 6a**). These results provide additional converging evidence that the brain representations of emotional valence are shared across people when the stimulus is less self-relevant, but they become idiosyncratic across people when the stimulus is highly self-relevant.

We then examined wherein the brain showed similar or distinct patterns of predictive weights between the valence models for high versus low self-relevance using a searchlight-based pattern similarity analysis method. As shown in **Fig. 6b**, the brain areas that showed low pattern similarity (in blue, Bayes Factor in favor of null hypothesis [BF_01_] > 6) were larger and more widely distributed across the whole brain than the brain areas that showed high pattern similarity (in red, Bayes Factor in favor of alternative hypothesis [BF_10_] > 6). Brain regions with low pattern similarity included cortical and subcortical regions within the default mode and limbic networks, such as the vmPFC, perigenual anterior cingulate cortex (pgACC), PCC, TP, hippocampus, and amygdala, and regions within the somatomotor network, such as supplementary motor area, right insula, and thalamus. Brain regions that showed high pattern similarity included the subgenual ACC (sgACC), left dorsal posterior insula (dpINS), and right dorsal lateral prefrontal cortex (dlPFC).

To summarize, these findings supported our hypothesis that the valence representations in the brain become more diverse and idiosyncratic across individuals as the stimuli become more self-relevant.

## Discussion

In this study, we replicated and significantly expanded our previous work using our newly developed Free Association Semantic Task (FAST) to assess the dynamic characteristics of the natural stream of thought^54^. Through an fMRI experiment (*n* = 62) combined with the FAST, we aimed to test whether we could predict individual differences in negative affectivity with dynamics of spontaneous thought, and whether we could identify and decode the brain representations and dynamics of phenomenological characteristics of spontaneous thought. Our main findings can be summarized as follows: (1) We developed a Markov chain-based predictive model of negative affectivity that generalized across multiple independent datasets. (2) Reflecting on one’s associative concepts strongly activated brain regions related to autobiographical memory, emotion, and internal and conceptual processing. (3) Predictive modeling of content dimension ratings of the self-generated concepts revealed that the brain representations of valence became more idiosyncratic as the level of self-relevance increased.

First, we successfully validated our new dynamic FAST, highlighting its great potential as a versatile thought sampling research tool for psychological assessment and neuroimaging studies. Here we showed that the FAST provided information about personally important thought topics and their semantic networks that could reveal each individual’s unique cognitive and phenomenological characteristics. The history of using free association as a psychological method to reveal one’s internal thoughts and emotional states goes back to the late 1800s^66-68^.

The method gained its increasing popularity since Sigmund Freud^92^, who used free association as the primary technique for his psychoanalysis, but modern psychology abandoned the method because of its questionable scientific validity and reliability. However, recent advances in computational tools and techniques provide an opportunity to revive the free association method with many potential use cases. For example, as shown in the current study, the dynamic modeling combined with machine learning has the potential to be used as an assessment tool for depression and anxiety in adjunct with self-report. Even in the absence of participant’s own self-report ratings, we previously showed in a behavioral FAST study that the affective dynamics of thought predicted individual differences in trait rumination^54^, a common symptom of mood and anxiety disorders. Thus, if we can implement an automated sentiment analyzer into the analysis pipeline, we can shorten the task time dramatically, providing a possibility to use the FAST as a web-or mobile-based monitoring tool for depression and anxiety. The FAST has a potential to be used for other clinical conditions, e.g., assessing spontaneous cognition in neurodegenerative disorders (e.g., Alzheimer’s disease^93^), thought disturbances in psychosis (e.g., derailment, flight of ideas, perseveration, etc.), or intrusive and repetitive thoughts in anxiety, obsessive-compulsive, or post-traumatic stress disorders. Furthermore, the FAST fully embraces ideas from emerging trends in neuroimaging studies, such as naturalistic and personalized approaches to high-dimensional neuroimaging data that emphasize multivariate representations and network-level dynamics. Together, our study provides a new research tool that will be useful for both behavioral and neuroimaging studies, creating a new possibility of capturing psychological and neurobiological processes previously challenging to study.

Second, our task’s primary brain targets were regions related to autobiographical memory, self-referential and emotion processing, and the monitoring and modulation of visceral and autonomic activity, including the medial prefrontal cortex and the medial temporal lobe structures within the default mode and limbic systems. While participants were reflecting about the self-generated concepts, these brain regions (referred to as “meaning-related” in **Figs. 3b** and **3c**) showed a delayed but strong activation after the transient stimulus-driven activity within the early visual and attentional orienting networks. These distinct temporal patterns of brain activity suggest that participants paid attention to the stimulus at the beginning of the trial, but then turned their attention inward and initiated endogenous cognitive and affective processes to reflect on the stimulus’ personal semantic meaning from a first-person perspective. These findings have significant basic science implications for the dynamic interplay between perceptually-coupled and internally-guided (i.e. “imaginative”) thought—processes which are often assumed to be antagonistic, but which must work together when attaching personal meaning to external stimuli as we show here^41,94,95^.

Third, predictive modeling of content dimension ratings revealed that the brain representations of emotional valence became more idiosyncratic (i.e. person-specific) as the level of self-relevance increased. When we trained the population-level predictive models, which capitalized on multivariate fMRI pattern information conserved across individuals, we were able to predict only the self-relevance dimension ratings. The important predictors of the self-relevance model included multiple regions within the default mode and limbic system, including the MPFC, PCC, TPJ, TP, NAc, and hippocampus, and most of these brain regions have been implicated in self-referential, mentalizing, autobiographical memory, visceral monitoring, and autonomic regulation in the previous literature^31,49,81-85,96-100^. For the valence dimension, we failed to develop a well-performing prediction model of valence when all data were combined for the modeling, but when we used only the trials with low self-relevance, we were able to develop a population-level prediction model of valence. This valence-lowself model also showed significant generalization to two independent datasets that used exogenous emotional stimuli (i.e., IAPS pictures), suggesting the existence of the generalizable valence codes in the brain when the emotional stimuli were less self-relevant. However, individuals exhibited idiosyncratic brain representations of valence when stimuli were highly self-relevant.

The findings of the idiosyncratic valence representations induced by self-relevant stimuli have important implications for emotion research. First, the results highlight the importance of the choice of stimuli and tasks for the study of emotion. Suppose we only use exogenous stimuli to induce emotions, such as movie, music, or pictures generated or selected by researchers. In that case, we might not be able to fully capture the brain representations and mechanisms of endogenous affective experiences. For this reason, we call for future research to develop and employ experimental paradigms that focus more on self-generated and naturalistic stimuli to target endogenous and first-person experience of emotions. Second, our results suggest that the valence codes in the brain can be modulated by the stimulus types and contexts, supporting the “affective modes” hypothesis (a brain subsystem can have different valence codes depending on affective states and contexts), rather than the “affective module” hypothesis (a brain subsystem is dedicated to a single valence code)^101^. Our study showed that the self-relevance level could serve as a crucial affective context that can produce significant changes in affective modes in the brain. Third, though the affective mode change appeared to occur in multiple brain regions distributed across the whole brain, regions within the default mode network and the limbic system, such as the TPJ, hippocampus/amygdala, vmPFC, and TP, seem to play a central role in this mode change. This is likely because of their involvement in episodic and semantic memories and emotion processes^16,46,102,103^. Episodic memory-related brain regions may provide rich autobiographical and personal contextual information to the semantic representations of valence, producing idiosyncratic representations of valence—i.e., turning a simple representation of good and bad (i.e., valence) into more complicated, nuanced, and personally unique valence representations. In addition to the interaction between memory and emotion processes, visceral monitoring and autonomic modulation in these DMN and limbic regions may also play an important role in modulating valence representations in the brain^41,99,100^. These DMN and limbic regions have been proposed to provide a subject-centered reference frame by integrating various visceral inputs^104^, and thereby serve as a basis for the subjective experience of “self-reletedness”^99^ (or “mineness”^105^). The importance of these regions in processing self-generated concepts was further supported by the supplementary analyses shown in **Supplementary Figs. 8-11**, but it still needs to be further examined in future studies (e.g., using the semantic encoding model^63,106^ or state space modeling^107^).

There are some considerations and limitations in the current study. First, some participants reported that the task was difficult to perform. For example, some participants became nervous about the MR scanner environment, and thus they felt difficulty engaging in free association with verbal report. Some participants reported that the 2.5-second interval for concept generation was too short even though it was allowed to repeat the previous response. Fortunately, our behavioral predictive model showed good generalizability across different experimental parameters, including the time limit for concept generation and response modality (e.g., the web-based FAST collected the response through typing with a 7-second interval). Thus, future studies could provide a longer time interval for the concept generation. Another factor that can influence task difficulty is verbal fluency. Although the verbal fluency scores did not show significant correlations with the model prediction or input features (see **Supplementary Table 5**), it may be because our participants were mostly college students with similar levels of verbal fluency. Therefore, to generalize our findings to a general population, future studies should further investigate the influences of experimental parameters and verbal fluency on task performance and difficulty. Second, though we analyzed the fMRI data from the concept reflection period to minimize the issues related to motion confounds, the brain dynamics and activity patterns during the actual free association (i.e., the concept generation period) could be different from the concept reflection period. To address this issue, we will need creative methods to mitigate motion confounds during free association, e.g., using a customized head mold^108^ (but also see ref. ^109^) or a silent speech interface^110^. Third, future studies should test different content dimensions, given that our choices on the dimension scales are somewhat arbitrary. To keep the whole experiment session including the post-scan survey within a reasonable time (< 4 hours), we had to choose only a few numbers of content dimensions (though we selected the current dimensions based on previous work^28^).

Overall, the current study opens up a new possibility of a quantitative assessment of the spontaneous thought dynamics and their brain representations. Our behavioral task and predictive models have the potential to be clinically useful because they can provide rich information about an individual’s cognitive and conceptual dynamic signatures^111^. Furthermore, our findings suggest that neural representations of affective processes become more idiosyncratic when self-relevant stimuli are used to induce emotions, highlighting the importance of targeting endogenous cognitive and affective processes in the study of emotion. Together, our study leads us to a deeper understanding of how self-relevance modulates the affective representations in the brain—in other words, what happens in our brain *when the self comes to a wandering mind*.

## Methods

### Participants

For the FAST-fMRI study, 63 healthy, right-handed participants participated (age = 23.0 ± 2.5 years [mean ± SD], 30 females). The preliminary eligibility of participants was determined through an online screening questionnaire. We did not include participants with psychiatric, neurological, systemic disorders or MRI contraindications. After the experiment, we excluded behavioral (and fMRI) data from one participant who generated too few responses. Thus, we used data from 62 participants in the behavioral data analysis. We also had to exclude one participant’s fMRI data due to the insufficient MRI coverage. Thus, we used data from 61 participants in the fMRI analysis. For the re-test session, 30 participants (age = 22.8 ± 2.3 years, 15 females) re-visited the experiment about seven weeks (mean = 51.0 ± 16.8 days) after their first visit. For the web-based FAST study that was used for an independent test data of the Markov chain analysis, 117 participants (age = 22.6 ± 2.6 years, 56 females) completed the web-based behavioral FAST experiment in the behavioral experimental room. Among them, 49 participants (age = 23.0 ± 3.1 years, 24 females) revisited and conducted the second session. Their revisits were about seven weeks (mean = 54.2 ± 8.5 days) after their first visits.

We recruited participants from Suwon area in South Korea, and the experiments were conducted at the Center for Neuroscience Imaging Research, Sungkyunkwan University in Suwon, South Korea. The institutional review board of Sungkyunkwan University approved these studies. All participants provided written informed consent and were paid for their participation.

### Self-report questionnaires

To assess individual differences in mental health and affective traits and states, all participants completed a battery of self-report questionnaires. In the fMRI study, we included the 20-item Positive and Negative Affect Schedule^112^ (PANAS, which had two subscales—positive affect and negative affect), the 20-item Center for Epidemiologic Studies Depression^113^ (CES-D), the 22-item Rumination Response Scale^114^ (RRS, which consisted of two subscales— brooding and depressive rumination), the 20-item trait version of the State-trait anxiety inventory^115^ (STAI-T) and the 30-item Mood and Anxiety Symptom Questionnaire-D30^116^ (MASQ-D30, which had three subscales—general distress, anhedonic depression, and anxiety arousal) in the battery. Participants completed their responses to the questionnaires prior to the fMRI scan.

In the web study, we included the 20-item PANAS, the 20-item CES-D, a subset of the RRS (9 items from the ‘depressive rumination’ sub-scale), and the 20-item STAI-T. These questionnaires overlapped with the fMRI study. In addition to these, we also included the Suicidal Ideation Questionnaire^117^ (SIQ; we only used 2 items, “I wished I were dead”, “I thought that life was not worth living”), the 3-item Loneliness Scale^118^ (LS), the 5-item Satisfaction With Life Scale^119^ (SWLS), and the 18-item Psychological Well-Being Scale^120^ (PWB, which had six subscales—autonomy, environmental mastery, personal growth, positive relations with others, purpose in life, and self-acceptance) in the battery. We used the Korean versions of these questionnaires that showed similar psychometric properties to the original questionnaires^121-127^.

### Free Association Semantic Task (FAST) for the fMRI experiment

The FAST for the fMRI experiment comprised the three parts—(1) concept generation, (2) concept reflection, and (3) post-scan survey. For the (1) concept generation phase, we asked participants to report a word or phrase that came to mind in response to the previous concept every 2.5 seconds starting from a given seed word inside the MRI scanner. The responses were collected through an MR-compatible microphone. Participants were asked to generate a total of 40 consecutive concepts for each seed word, and we used four seed words for four runs total. We made the number of associations for each seed word much longer than our previous study^54^, in which we collected only 10 consecutive concepts, in order to obtain a larger number of personal concepts. The four seed words were “family”, “tear”, “mirror”, and “abuse” for the first session and “love”, “fantasy”, “heart”, and “pain” for the re-test session. The selection of the seed words was based on a pre-survey of valence and self-relevance with a large set of candidate words. We selected these seed words to make them evenly distributed on the valence and self-relevance dimensions. The orders of the seed words were fully randomized across participants.

During the (2) concept reflection phase, we showed participants two consecutive concepts generated by themselves in sequence. We made the second concept bigger than the first one to make it clear for the second concept to be the target word. Then, we asked participants to think about the target concept and personal context relevant to the association between the two concepts for 15 seconds. The stimuli remained on the screen for the whole duration (i.e., 15 seconds). We tried to give them enough time to think about their personal context, such as memory that gave the concept a personal meaning. Between the trials, we showed a fixation cross with a jittered duration between 3 and 9 seconds (i.e., inter-stimulus interval “baseline”). Intermittently, we showed fourteen emotion words on the screen after the word presentation and asked participants to select one emotion descriptor closest to their current feeling. The emotion words were joy, distress, hope, fear, satisfaction, disappointment, pride, embarrassed, remorse, gratitude, anger, love, hate, and neutral. We conducted this emotion rating five times per run and collected a total of 160 self-generated words per participant. We used these emotion rating data to validate the post-scan survey results (**Supplementary Fig. 2**).

After the fMRI scan, participants completed a (3) post-scan survey on the 160 self-generated concepts in the behavioral experiment room. We showed the self-generated concepts again and asked participants to rate them on multi-dimensional content dimensions^28^. The content dimensions evaluated emotional valence (how much positive or negative feelings does the concept evoke?), self-relevance (how much is the concept relevant to yourself?), time (which time point is most relevant to the concept, ranging from the past to the future?), vividness (how much vivid imagery does the concept induce?), and safety-threat (does the concept give rise to the feeling of safety or threat?). Like the concept reflection task, we showed two consecutive concepts in sequence with five content dimension questions and asked participants to answer the questions about the target concept (i.e., the second one). We used the visual analog scale. The valence, time, and safety-threat ratings were coded between -1 (negative, past, threat, respectively) and 1 (positive, future, safety), and 0 indicated neutral or present. The self-relevance and vividness ratings were coded between 0 (not self-relevant, not vivid) and 1 (highly self-related, highly vivid).

### FAST for the web experiment

The web-based FAST consisted of the concept generation and concept survey tasks. The concept generation task was similar to that of the fMRI experiment, but this time we provided 10 seconds for the concept generation because typing usually took longer than speaking. The seed words for the first and second sessions were the same as those of the fMRI experiment. After the concept generation task, participants completed the concept survey on three content dimensions—valence, self-relevance, and time. We chose these three dimensions based on the principal component analysis (PCA) results shown in **Supplementary Fig. 3a**. The PCA results suggested that the valence and safety-threat dimensions were highly correlated, and the self-relevance and vividness were also highly correlated.

### Markov chain-based predictive modeling of negative affectivity

To build predictive models of general negative affectivity, we used features obtained from a Markov chain analysis on the content dimensions. First, we divided the dimension scores into multiple discrete states. For the valence, time, and safety-threat dimensions, which ranged from -1 to 1, we divided them into three discrete states using -0.33 and 0.33 as the boundaries for defining discrete states (i.e., -1 to -0.33, -0.33 to 0.33, and 0.33 to 1; for valence, the three discrete states were negative/neutral/positive, for time, past/present/future, for safety-threat, threatening/neutral/safe). The self-relevance and vividness dimensions, ranging from 0 to 1, were divided into two discrete states (0 to 0.5 and 0.5 to 1, which corresponded to low and high for both dimensions, respectively).

We then calculated the state transition and steady state probabilities for each dimension and for each participant. The state transition probability refers to the probability of making transitions from one to another discrete state on each dimension. The steady state probability refers to the probability of converging to one state when the transitions were sufficiently repeated^71^. We obtained the steady state probability by multiplying the transition probability matrix by itself 10,000 times, which was always converged to one probability vector. In addition to these dynamic features from the Markov chain analysis, we also used each content dimension’s mean and variance as input features. Together, we used nine (= 3 × 3) transition probability values and three steady state probabilities for the valence, time, safety-threat dimensions, four (= 2 × 2) transition probabilities and two steady state probabilities the self-relevance and vividness dimension, and mean and variance of all dimensions, creating a total of 58 input features (i.e., predictor variables). Most of these features showed a good level of consistency across different sets of seed words and different time points (**Supplementary Table 1**).

For the outcome variable, we conducted factor analyses to calculate general negative affectivity scores from a combination of self-report questionnaires. As an input for the factor analyses, we used z-scored subscale scores because all questionnaires used different scales. We used a target oblique rotation for a two-factor model, given that we had *a priori* knowledge about what each questionnaire measures between positive versus negative affectivity. The general negative affectivity factor consisted of questionnaires related to negative affect, general distress, anxiety, and depression (**Supplementary Table 2**), which was then used as the outcome variable for the predictive modeling.

With these predictor and outcome variables, we developed a predictive model of the negative affectivity using least absolute shrinkage and selection operator (LASSO) regression^72^. We used the first session data of the fMRI study (*n* = 62) as a training dataset and tested the developed model on the second session data of the fMRI study (*n* = 30) and two session (test and re-test) datasets of the web study (*n* = 117 and 49, respectively). We determined the number of the predictor variables for the final model based on the leave-one-participant-out cross-validation performance in the training set with the LASSO regularization.

To test the prediction model on independent test datasets, we calculated the predicted levels of general negative affectivity using a dot product between the Markov chain features calculated from new datasets and the model weights. To evaluate the model performance, we used robust correlation between the actual and predicted levels of general negative affectivity. We did not use mean squared error or R-squared to evaluate model performance as recommended in ref. ^76^ given that the scale of outcome variables were different between the training and test datasets because they used the different sets of self-report questionnaire.

### fMRI data acquisition and preprocessing

We collected MRI data using a 3T Siemens Prisma scanner at Sungkyunkwan University. We acquired high-resolution T1-weighted structural images and functional EPI images with TR = 460 ms, TE = 27.2 ms, multiband acceleration factor = 8, field of view = 220 mm, 82×82 matrix, 2.7×2.7×2.7 mm^3^ voxels, 56 interleaved slices, number of volumes = 2608. Stimulus presentation and behavioral data acquisition were controlled using MATLAB (Mathworks®, Natick, MA) and Psychtoolbox (http://psychtoolbox.org/)

Data preprocessing were performed with SPM12 (Wellcome Trust Centre for Neuroimaging) and FSL (the Oxford Centre for Functional MRI of the Brain). For structural T1-weighted images, we co-registered the T1 images to the functional image using the first single band reference (SBRef) image and segmented and normalized to the MNI space. For the functional EPI images, we removed the initial volumes (20 images) of fMRI data to allow for image intensity stabilization. We also identified outliers for each image and all slices based on Mahalanobis distances and the root mean square of successive difference to remove intermittent gradient and severe motion-related artifacts that are present to some degree in all fMRI data. For the Mahalanobis distance-based outlier detection, we computed Mahalanobis distances for the matrix of concatenated slice-wise mean and standard deviation values by volumes across time. Then we identified the images that exceed 10 mean absolute deviations (MADs) based on moving averages with full width at half maximum (FWHM) 20 images kernel as outliers. With the root mean square of successive difference across volumes, images that exceeded 3 standard deviations from the global mean were identified as outliers. Each time-point identified as outliers by either outlier detection method was included as nuisance covariates.

We then conducted 1) motion-correction (realignment) using the SBRef image as a reference, 2) distortion-correction using FSL’s topup function, 3) normalization to the MNI space using the parameters from the T1 normalization with the interpolation to 2×2×2 mm^3^ voxels, and 4) smoothing with a 5-mm FWHM Gaussian kernel. Since data from two participants showed poorer quality images after the distortion correction, we used distortion-uncorrected images for these two participants.

### fMRI single trial analysis

We used the single trial, or ‘single-epoch,’ analysis approach. We estimated single-trial response magnitudes for each brain voxel using a GLM design matrix with separate regressors for each trial, as in the ‘beta series’ approach^128^. We constructed each trial regressor for the concept reflection duration with a boxcar convolved with SPM12’s canonical hemodynamic response function. We also included a regressor for intermittent emotion ratings for each run. We concatenated four runs’ data during the concept reflection task for each participant and added the run intercepts. In the design matrix, we also included the nuisance covariates, including (i) ‘dummy’ coding regressors for each run (an intercept for each run), (ii) linear drift across time within each run, (iii) 24 head motion parameters (6 movement parameters including x, y, z, roll, pitch, and yaw, their mean-centered squares, their derivatives, and squared derivative) for each run, (iv) indicator vectors for outlier time points, and (v) five principal components of white matter and cerebrospinal fluid signal. With this design matrix, we ran the first level analysis using SPM12 with a high-pass filter of 180 seconds. Since single trial estimates could be strongly affected by acquisition artifacts that occur during that trial (for example, sudden motion, scanner pulse artifacts, etc.), we calculated trial-by-trial variance inflation factors (VIFs, a measure of design-induced uncertainty due to collinearity with nuisance regressors) using the design matrix, and any trials with VIFs that exceeded 3 were excluded from the following analyses. The average number of trials excluded due to high VIFs was 2.902 with the standard deviation of 3.081.

### Large-scale functional network overlap analysis

The radial plots in **Figs. 3-4** and **6** show the relative proportions of the number of overlapping voxels between the thresholded map and each network (or region) given the total number of voxels within each network (or region). We used the Buckner group’s parcellations to define large-scale functional brain networks, including 7 networks within the cerebral cortex^129^, cerebellum^130^, and basal ganglia^131^. We also added thalamus, hippocampus and amygdala from the SPM anatomy toolbox^132^ and the brainstem region, as shown in **Supplementary Fig. 5**.

### Clustering analysis using the finite impulse response model

For the clustering analysis based on the temporal patterns of brain activity, we estimated the TR-level brain activity patterns using the finite impulse response model (**Figs. 3b-c**). We modeled a total of 35 TRs (= 16.1 seconds) from the concept reflection trial onset. We used the thresholded univariate contrast map (at *p* < .05, Bonferroni correction) for the concept reflection duration as a mask. We then conducted *k*-means clustering on the voxels within the mask based on the temporal patterns of brain activity across 35 TRs. We selected the number of clusters *k* that maximized the Silhouette value of the clustering solution using “evalclusters.m” function included in the MATLAB statistical toolbox. To show the time course of the cluster brain activity as in **Fig. 3b**, we plotted the averaged beta weights within each cluster at each time point.

### Predictive modeling of content dimensions

To develop multivariate predictive models of each content dimension, we first divided the trials into four levels and averaged the dimension ratings and fMRI data for each level, creating the quartile training data for each participant. For each dimension, a total of 244 images (4 [four images per participant] × 61 [# of participants]) were created for model training. Some dimensions had less than 244 images due to the skewed distribution of some participants’ ratings or the removal of some trials with high VIFs. We used principal component regression (PCR) to train multivariate pattern-based predictive models. To obtain unbiased estimates of model performance, we used two different types of cross validation methods. The first one was the leave-one-subject-out cross-validation, in which we derived a predictive map from all participants’ data except one participant and used the hold-out participant’s data for the model testing. The other method was the random split cross-validation^77^, in which we randomly chose 20% of participants’ data as the hold-out data for each iteration. We then used 80% of randomly selected participants’ data to derive a predictive map and used the 20% hold-out data for the model testing for each iteration. We repeated this procedure 50 times. We evaluated model performance with 1) averaged within-participant correlation between actual and predicted ratings and 2) averaged within-participant mean squared error. To test whether the mean within-participant prediction-outcome correlation was significantly larger than zero, we conducted bootstrap tests with participant-level prediction-outcome correlation values with 10,000 iterations. For the predictive model of valence only for low self-relevance trials (i.e., the valence-lowself model; **Fig. 4b**), we divided the data into trials with high versus low self-relevance scores using 0.5 as a threshold before making the quartile data. After that, all the model training and testing steps were the same as above.

To help the feature-level interpretation of the predictive models, we conducted bootstrap tests with 10,000 iterations. For each iteration, we randomly sampled participants with replacement and trained a PCR model using the resampled dataset. Based on the sampling distribution of bootstrapped predictive weights, we identified features that consistently contributed to the prediction with *p*-values. For display as in **Fig. 4**, we thresholded the map with the false discovery rate (FDR) *q* < 0.05 and pruned the results using two additional more liberal thresholds, uncorrected voxel-wise *p* < .01 and *p* < .05, two-tailed.

To test our valence-lowself model on an independent study dataset, we obtained predicted ratings using a dot product of each vectorized test image data with model weights. To evaluate the prediction performance, we used averaged within-participant correlation between actual and predicted ratings. We did not use the mean squared error because the units were different between the training and testing datasets. Similarly, to test the Picture-Induced Negative Emotion Signature (PINES) on our data, we calculated pattern expression values with a dot product between the test data and the PINES weights and used averaged within-participant prediction-outcome correlation as the evaluation measure.

### Idiographic predictive modeling of valence

To quantify the between-subject variability of predictive weights for data with high versus low self-relevance, we used the idiographic predictive modeling approach. First, we divided trials into high and low self-relevance groups using 0.5 as a cutoff score. For each participant, we trained two predictive models of valence with these two groups of trials separately using PCR with 5-fold cross-validation.

We also conducted a searchlight-based pattern similarity analysis to identify brain areas that showed similar or distinct patterns of predictive weights between two types of valence prediction models. For this, we created a searchlight with a radius of 5 voxels and scanned it throughout the whole brain with a step size of 4 voxels. We then calculated correlation coefficients between two models’ predictive weights within each searchlight and transformed *r* to *z* using Fisher’s transformation. We added the *z*-values to cubes which had one side of 4 voxels and were located at the center of each searchlight across the whole-brain with no overlapping voxels between cubes. We smoothed the map with a 3mm FWHM Gaussian kernel and performed one-sample *t*-test, treating participants as a random effect. We also calculated the JZS Bayes factors using the method proposed in ref. ^133^.

## Data Availability

The predictive models and the data to generate the main figures will become available at https://cocoanlab.github.io/fast_fmri upon publication.

## Code Availability

Codes for generating main figures will be available at https://cocoanlab.github.io/fast_fmri upon publication. In-house MATLAB codes for fMRI data analyses (e.g., preprocessing and predictive modeling) are available at https://github.com/canlab/CanlabCore and https://github.com/cocoanlab/cocoanCORE.

## Acknowledgements

We thank Minie Jung, Jinwon Park, and Seok Ho Song for help with conducting experiments. This work was supported by IBS-R015-D1 (Institute for Basic Science; to C.-W.W.), 2019R1C1C1004512, 2021M3A9E4080780, and 2021M3E5D2A0102249311 (National Research Foundation of Korea; to C.-W.W.), 2E30410-20-085 (KIST Institutional Program; to C.-W.W.).

## Author Contributions

B.K., J.R.A., and C.-W.W. conceived and designed the fMRI experiment. B.K. and C.-W.W. conducted the experiment, analyzed the data, interpreted the results, and wrote the manuscript. B.K., J.R.A., and C.-W.W. revised the manuscript. C.-W.W. provided the supervision. B.K., J.R.A., and C.-W.W. contributed to the FAST-fMRI study. B.K., J.H., E.L., and C.-W.W. contributed to the FAST-web study.

## Supplementary Information

Supplementary Figures 1-11

Supplementary Tables 1-5

