## Supplementary information for "When self comes to a wandering mind: Brain representations and dynamics of self-generated concepts in spontaneous thought"

**This PDF file includes:**

1. Supplementary Figures 1-11
3. Supplementary Tables 1-5

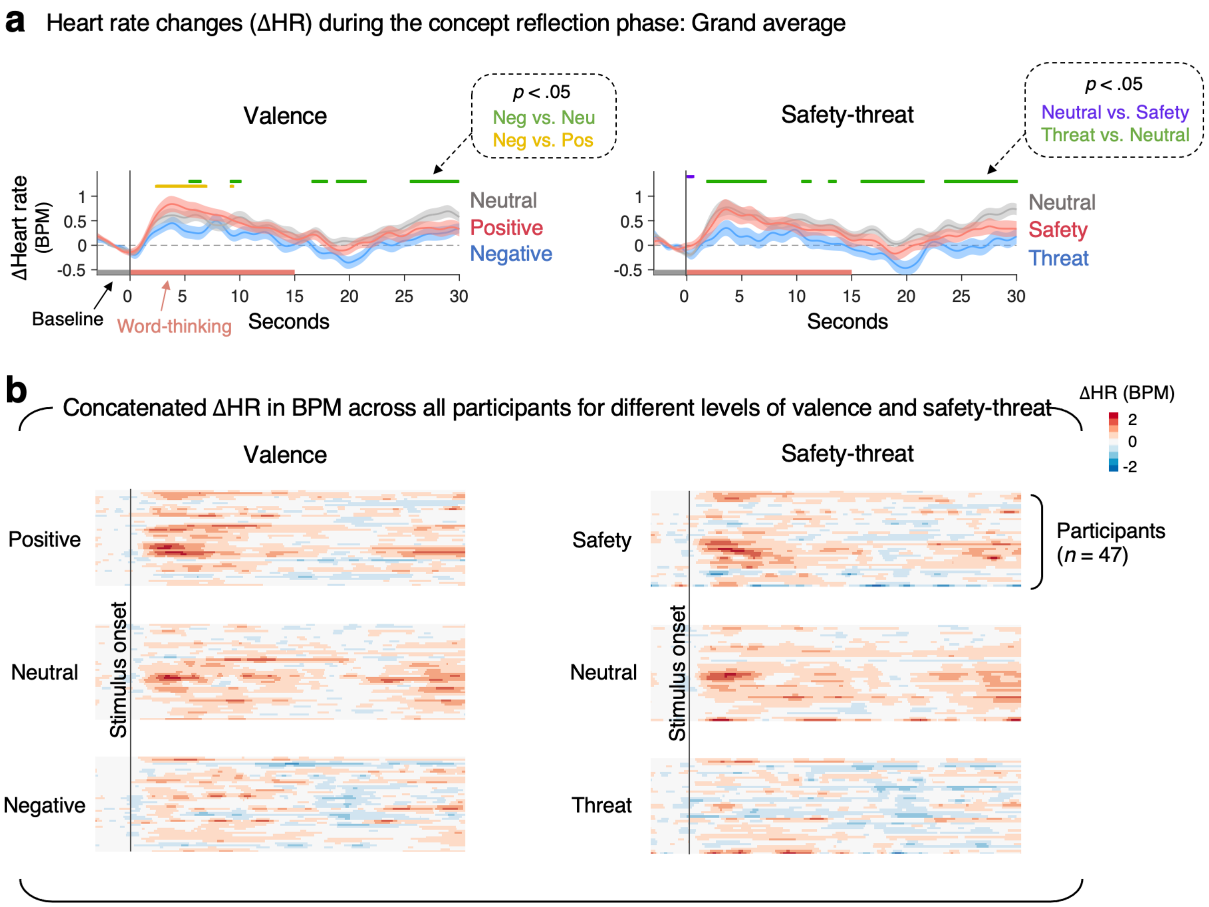


**Supplementary Figure 1. Valence-induced heart rate changes during the concept reflection phase. a**, The plots show the time-course of grand-average heart rate changes (ΔHR; in beats per minute [BPM]) during the concept reflection phase. Shading represents standard error of the mean (s.e.m.) across participants. **b,** The heat maps show the concatenated grand-average ΔHR across all participants for different levels of valence and safety-threat. The red color means a faster heart rate than baseline and the blue color means a slower heart rate than baseline.

**Main finding:** There were multiple time-points that showed significant decreases in heart rate for the negative vs. neutral (or positive) and threat vs. neutral contrasts.

**Methods.** Electrocardiogram (ECG) activity was recorded using MR-compatible electrodes (Biopac systems, Goleta, CA) placed on the right and left clavicle and lower left abdomen area. We analyzed the ECG data from the first session of the fMRI experiment ($n$ = 62). We had to discard 15 participants' data—5 participants due to the issues related to the recording setup and 10 participants due to abnormal BPM ranges due to MR-related noise. Thus, we used data from 47 participants in this analysis. The ECG data were sampled at 2000Hz during the scan. To remove the MR-induced noise from the ECG data, we conducted the band-pass filtering (0.6 ~ 10 Hz) and also used a comb filter with the multiple of a reciprocal number of TR (1/0.46 Hz). We then used the PhysIO Toolbox[1] (https://www.nitrc.org/projects/physio/) to find peaks and calculated the inter-beat interval (IBI). The IBI data were down-sampled (25 Hz) and low-pass filtered (0.5 Hz). Then we calculated beats per minute (BPM) by dividing 60 seconds by IBI (60/IBI). To examine the heart rate changes induced by the levels of valence and safety-threat scores, we divided the data into three groups—positive, neutral, and negative for valence, and safety, neutral, and threat for safety-threat—using 0.33 and -0.33 as the boundaries for defining discrete states. We grand-averaged the HR data using 3 seconds before the onset as the baseline and 30 seconds after the onset as an epoch. We conducted paired *t*-test for each time point, and the green, yellow, and purple dots (or lines) in the plots in **a** show the time points that yielded the significant *t*-test results, uncorrected $p < 0.05$, two-tailed, paired *t*-test, $n = 47$.

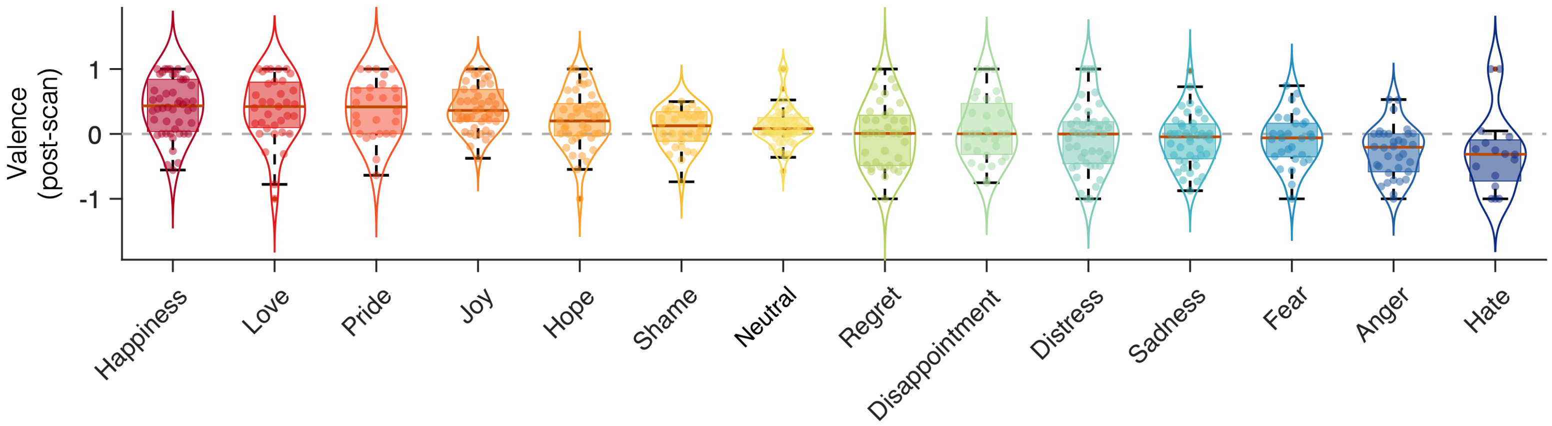


**Supplementary Figure 2. The relationship between the in-scanner emotion ratings and post-scan valence ratings.** The plot shows the distribution of the valence ratings (from the post-scan survey) for different emotion categories of self-generated concepts (*n* = 62). The emotion category data were from intermittent emotion ratings during the concept reflection task inside the scanner. For emotion ratings, we intermittently displayed 14 emotion words on the screen after 15 seconds of the concept presentation and asked participants to select one emotion descriptor closest to their current emotion. We obtained the emotion ratings five times per run. The box was bounded by the first and third quartiles, and the whiskers stretched to the greatest and lowest values within median ± 1.5 interquartile range.

**Main finding:** The post-scan valence scores were largely consistent with the in-scanner emotion ratings obtained during the fMRI experiment.

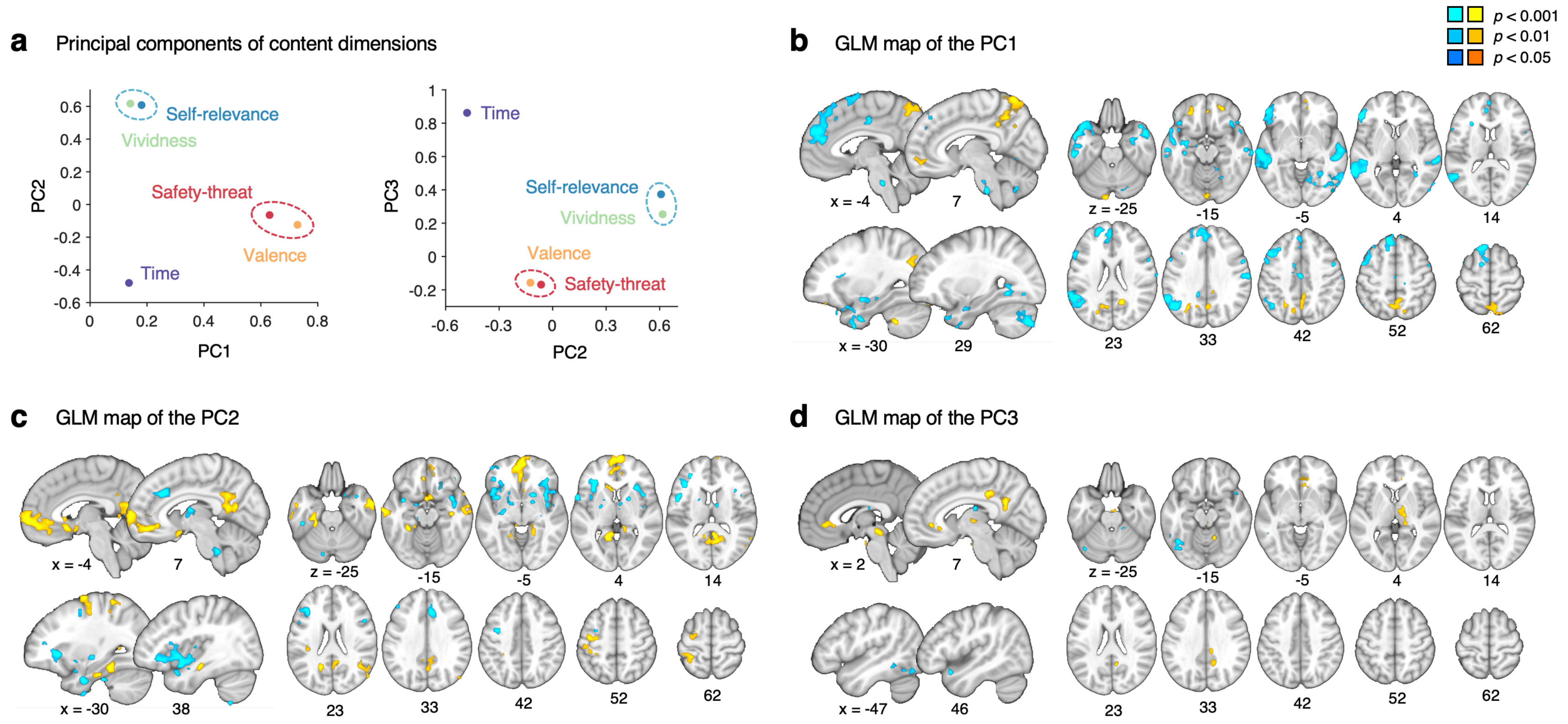


**Supplementary Figure 3. Principal components of the content dimensions and their univariate general linear model maps. a,** We conducted the principal component analysis on the five content dimensions ($n = 61$) and plotted three principal components (PCs), which explained 89.4% of the total variance. The valence and safety-threat dimensions showed high loadings on the first PC, while the self-relevance and vividness dimensions showed high loadings on the second PC. The time dimension showed a high loading on the third PC. **b-d**, The general linear model (GLM) results for the three principal components. To identify brain regions correlated with the principal component scores, we regressed the single trial images on the three principal component scores for each participant. We then performed a one-sample *t*-test on the 61 beta images (one beta map per participant) and thresholded the map with uncorrected $P < .001$, two-tailed, and pruned the results using two additional more liberal thresholds, $p < .01$ and $p < .05$, two-tailed.

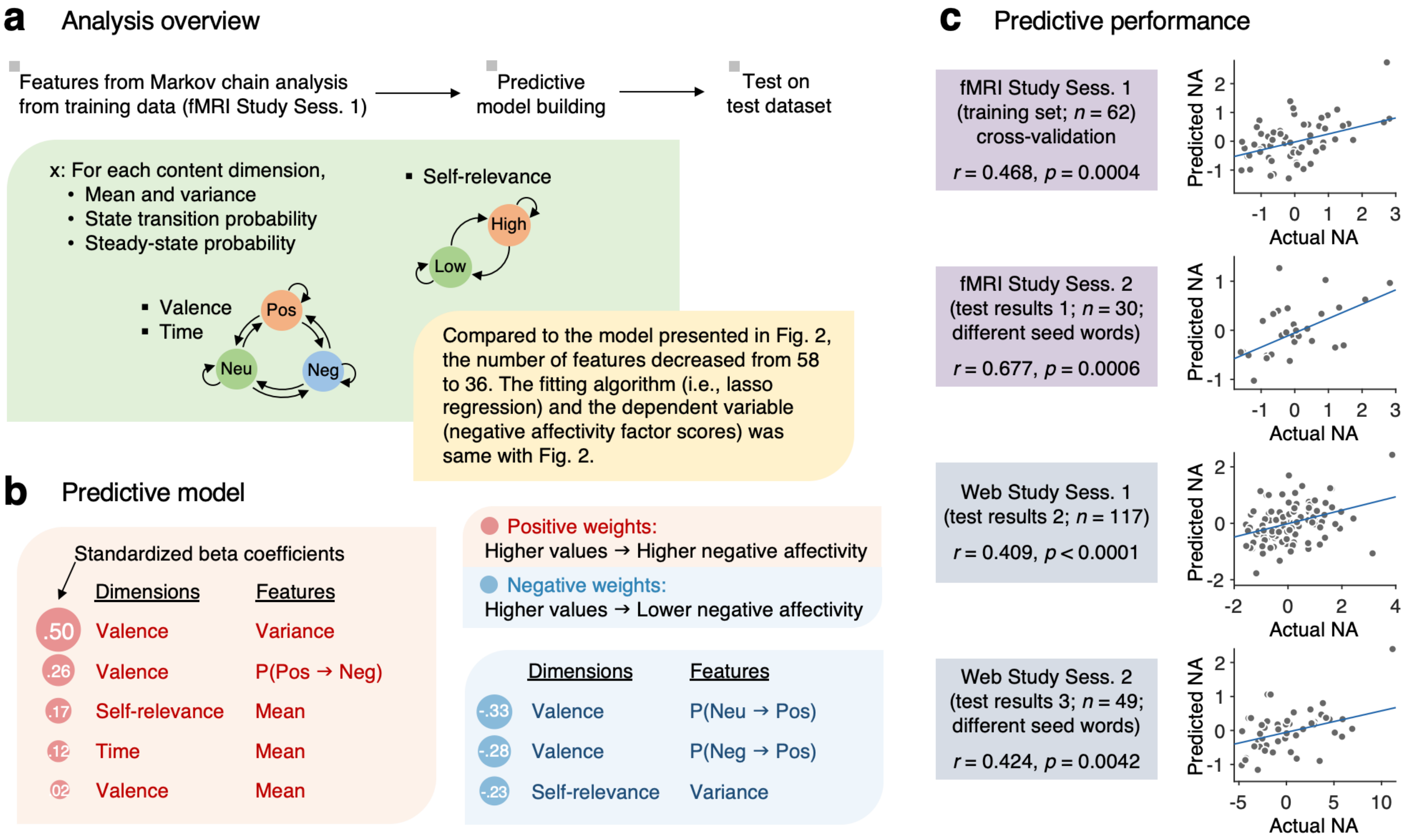


**Supplementary Figure 4. Markov chain-based predictive model of negative affectivity only based on valence, self-relevance, and time (VST model). a**, We trained an additional Markov chain-based predictive model of general negative affectivity only with the valence, self-relevance, and time dimensions to test whether these three dimensions were enough to predict the level of general negative affectivity. Other analysis procedure was identical to the main Markov chain-based predictive model with the full five content dimensions (**Fig. 2**). The total number of input features decreased from 58 to 36. **b,** A total of 8 features were selected. All the features except for the "valence-mean" overlapped with the selected features in the original model. **c**, VST model performance. From top to bottom, the plots show 1) the leave-one-participant-out cross-validated prediction results within the training dataset ($n$ = 62, first session of the fMRI study), and three independent test results on 2) the second session re-test data of the fMRI study with different seed words ($n$ = 30), 3) the first session data of the FAST-web study ($n$ = 117), and 4) the second session re-test data of the FAST-web study with different seed words ($n$ = 49). The actual versus predicted negative affectivity factor scores are shown in the plots. Each dot represents each participant. We evaluated the model performance with robust correlation between the actual and predicted levels of general negative affectivity.

**Main finding:** The VST model also showed significant predictions across four datasets, and seven out of eight final features of the VST model overlapped with the original full model (**Fig. 2c**).

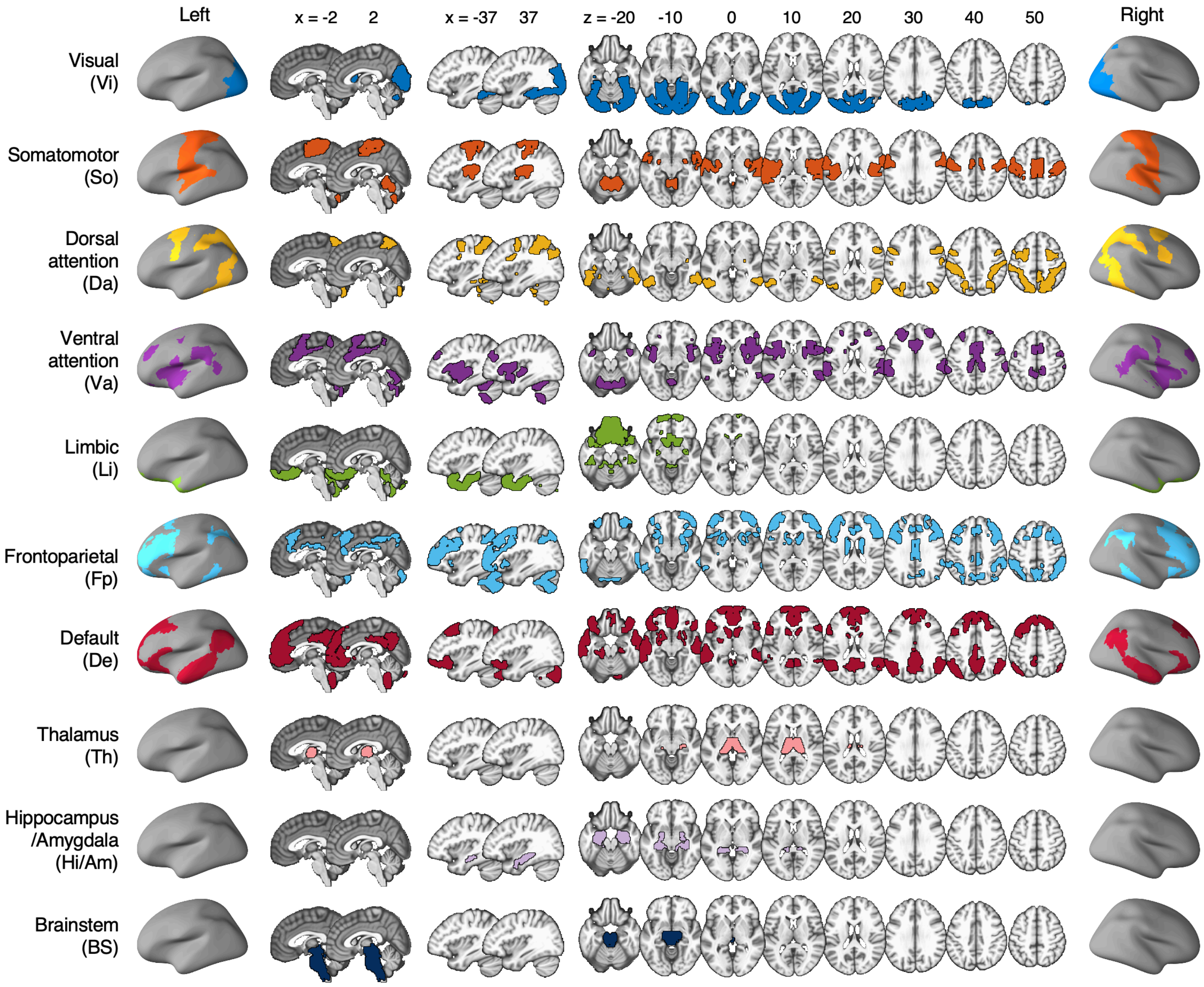


**Supplementary Figure 5. Large-scale functional networks and regions for the radial network plots.** To make the radial plots in the main figures (**Figs. 3, 4,** and **6**)**,** we used the Buckner group's parcellations to define large-scale functional brain networks, including 7 networks within the cerebral cortex[2], cerebellum[3], and basal ganglia[4]. We also added thalamus, hippocampus and amygdala from the SPM anatomy toolbox[5] and the brainstem region.

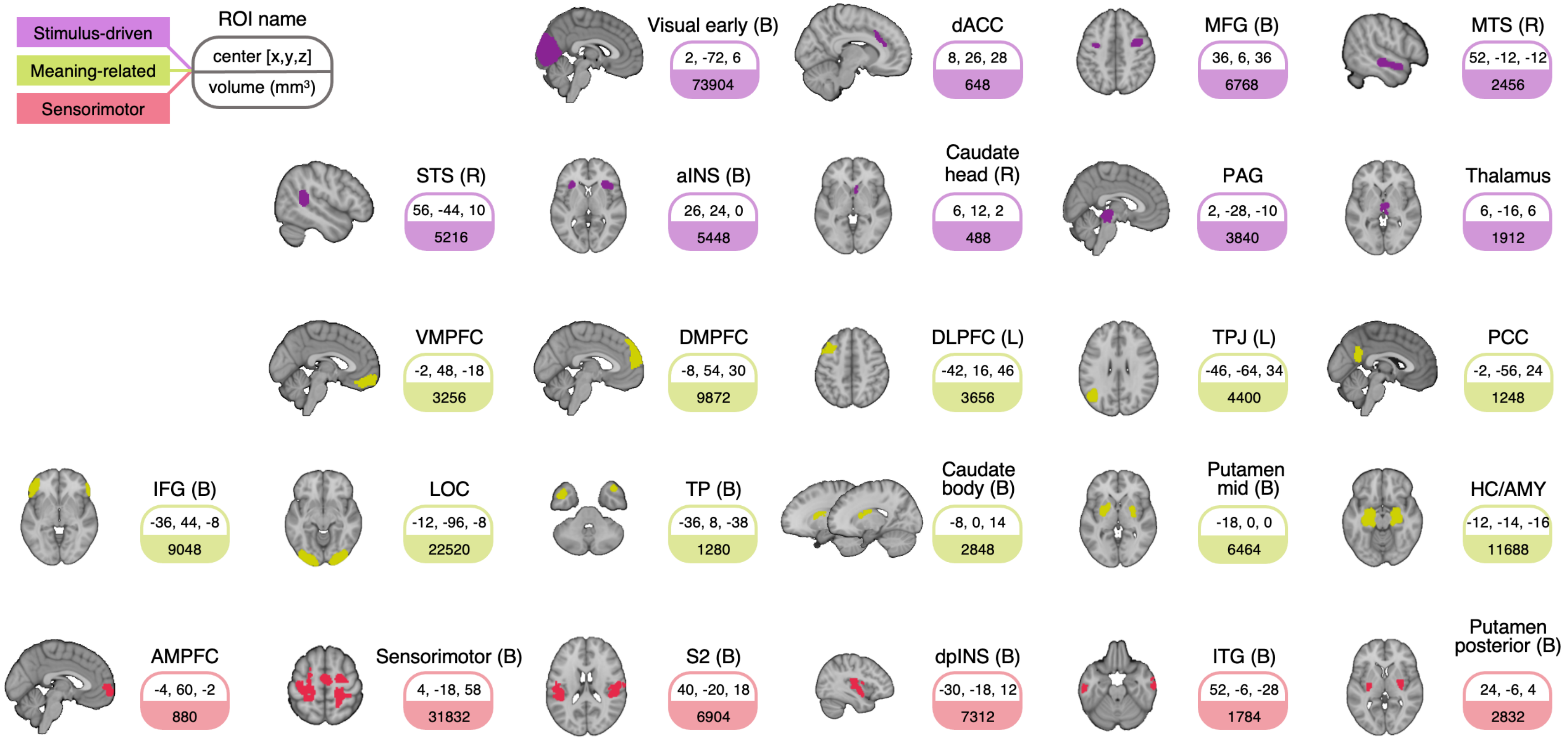


**Supplementary Figure 6. 26 main regions-of-interest (ROIs) from the basic contrast map of concept reflection.** dACC, dorsal anterior cingulate cortex; MFG (B), bilateral middle frontal gyrus; MTS (R), right middle temporal sulcus; STS (R), right superior temporal sulcus; aINS (B), bilateral anterior insula; PAG, periaqueductal gray; VMPFC, ventromedial prefrontal cortex; DMPFC, dorsomedial prefrontal cortex; DLPFC (L), left dorsolateral prefrontal cortex; TPJ (L), left temporal parietal junction; PCC, posterior cingulate cortex; IFG (B), bilateral inferior frontal gyrus; LOC, lateral occipital cortex; TP (B), bilateral temporal pole; HC/AMY, hippocampus/amygdala; AMPFC, anterior medial prefrontal cortex; S2 (B), bilateral secondary somatosensory cortex; dpINS (B), bilateral dorsal posterior insula; ITG (B), bilateral inferior temporal gyrus.

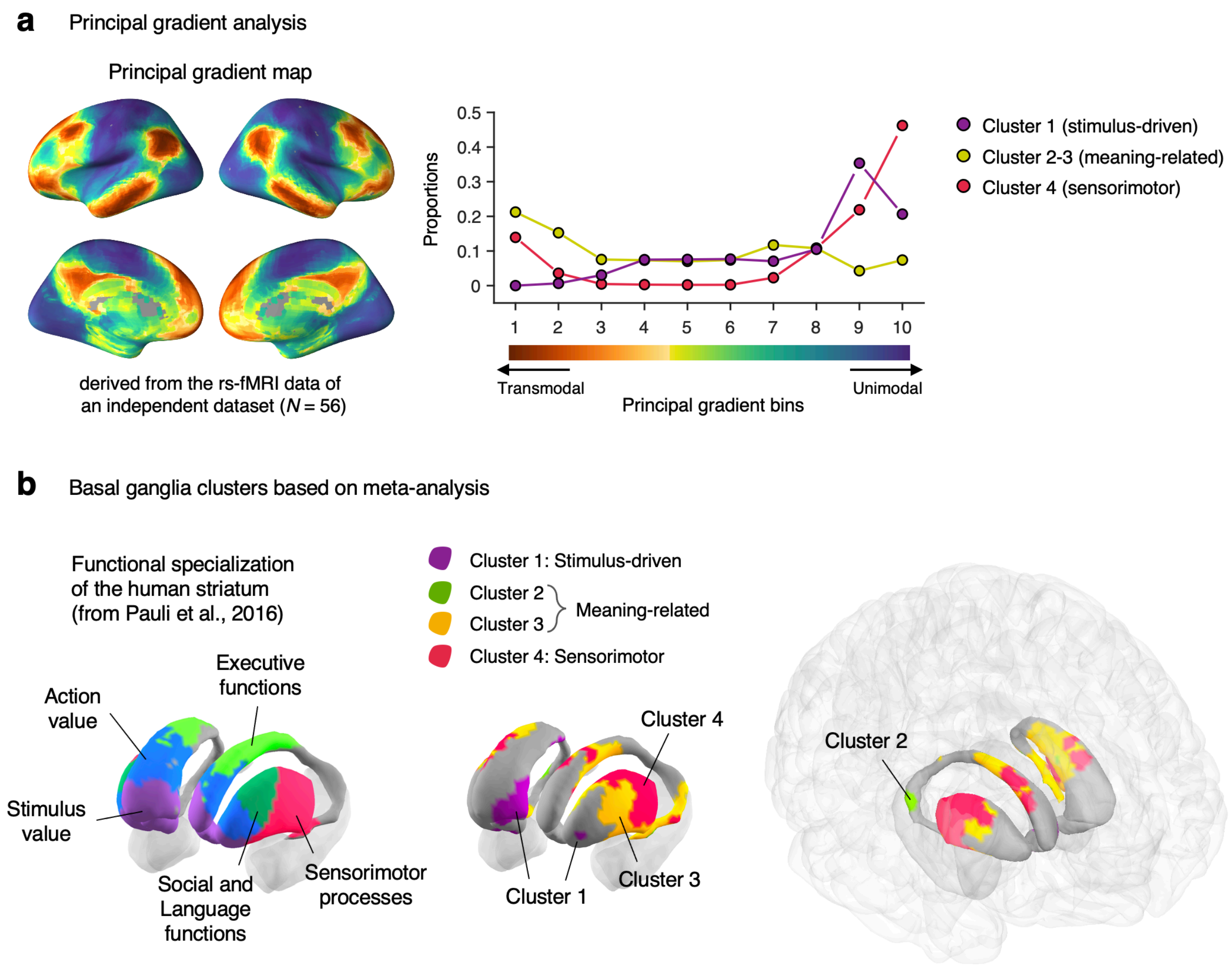


**Supplementary Figure 7. Neurobiological assessment of clustering results.** We assessed whether our clustering results and their naming (shown in **Fig. 3**) were neurobiologically and functionally meaningful rather than being arbitrary. In addition to the term-based decoding analysis reported in **Fig. 3c**, here we interpreted our clustering results with the principal gradient[6] and the meta-analytic basal ganglia parcellations[7]. **a**, The brain map on the left shows the principal gradient from the unimodal to trans-modal brain regions across the whole brain[6]. We re-calculated the principal gradient map using our own resting-state dataset (*n* = 56; 7-min resting scan) to create a volumetric principal gradient image and also to include the subcortical regions. The plot on the right shows the proportions of overlapping voxels between the 10-bin maps of the principal gradient and our three clusters. **b**, The basal ganglia map on the left shows the functional parcellations based on metaanalysis[7]. The figures in the middle and the right panels show our clustering results mapped on the basal ganglia.

**Main findings:** Our region clustering and their naming were largely consistent with the principal gradient in the cortex and the meta-analysis findings in the basal ganglia. For example, our clusters named “meaning-related” largely overlapped with the trans-modal end in the principal gradient of cortical hierarchy and the parts of the basal ganglia related to social, language, and executive functions. The “stimulus-driven” and “sensorimotor” clusters overlapped with the unimodal end of the cortical principal gradient and the basal ganglia parcellations for stimulus value and sensorimotor processes, respectively. These findings support that our clustering analysis resulted in neurobiologically meaningful clusters, providing a basis for further analyses of our data and functional interpretations of our findings.

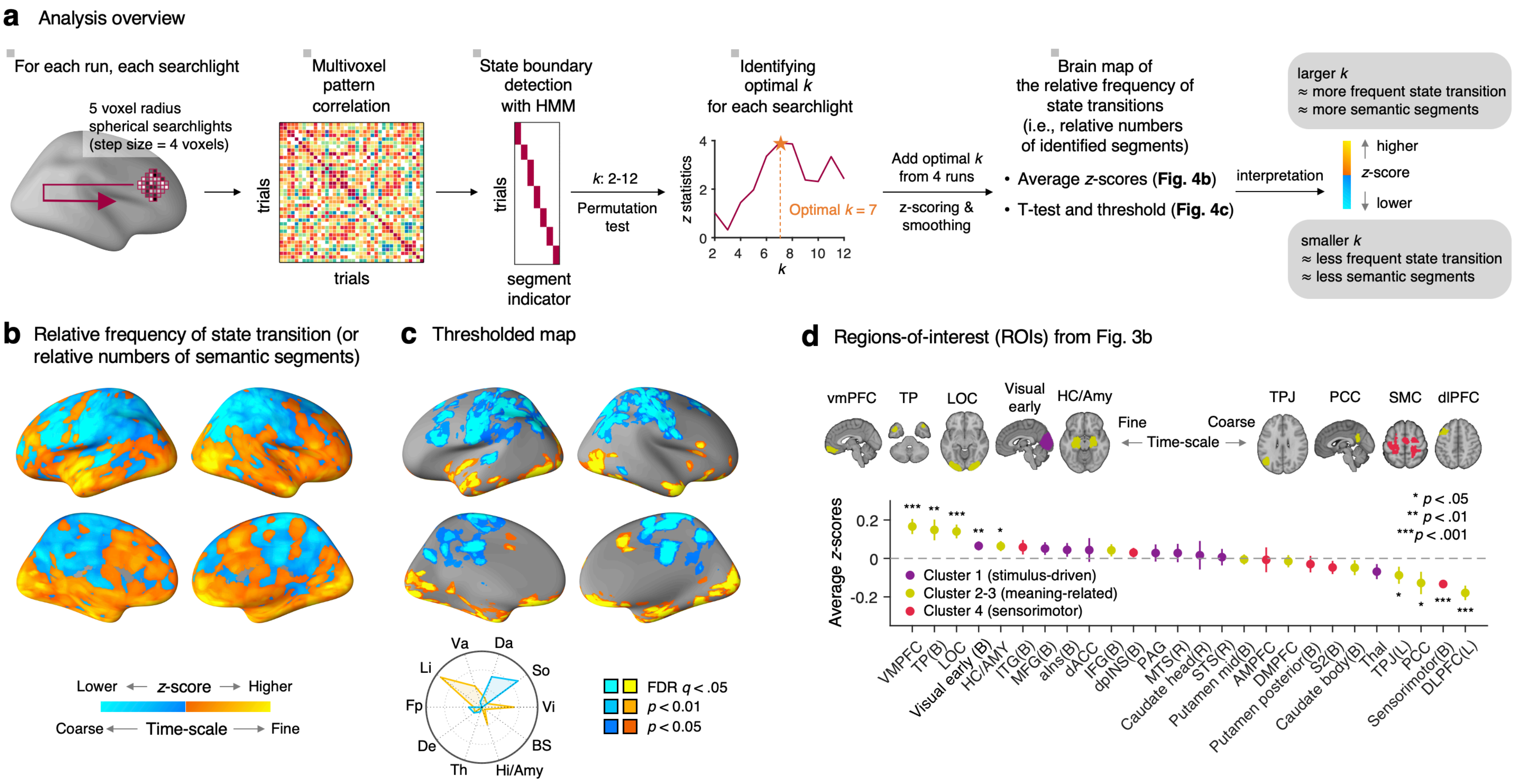


**Supplementary Figure 8. Relative frequency of state transition (or relative numbers of semantic segments) across the brain.** **a**, Analysis overview. To examine the relative frequency of state transition (or the relative numbers of semantic segments) of different regions based on their multivariate pattern information, we used a data-driven approach to detecting state boundaries with the Hidden Markov Model (HMM)[8]. For details about the analysis steps, please see below. **b**, Unthresholded group-level average of the state transition frequency. This map shows the group-averaged *z*-scores based on the total number ($k$) of state transition across four runs. The cool color indicates a smaller number of state transition, whereas the warm color indicates a larger number of state transition. **c**, Thresholded map of the relative frequency of state transition with FDR corrected $q < 0.05$, one-sample $t$-test, two-tailed. To better show the extent of the significant areas, we pruned the results using two additional more liberal thresholds, uncorrected voxel-wise $p < .01$ and $p < .05$, two-tailed. The radial plot shows the relative proportions of overlapping voxels between the thresholded map and large-scale networks. Va, ventral attention; Da, dorsal attention, So, somatomotor; Vi, visual; BS, brainstem; Hi/Am, hippocampus/amygdala; Th, thalamus; De, default; Fp, frontoparietal; Li, limbic. **d**, We conducted bootstrap tests for the 26 regions-of-interest (ROIs) obtained from the basic contrast map of the concept reflection-related brain activity. The plot shows the group-average $z$-scores with the standard error of the mean (s.e.m.). $^{*}p < .05$, $^{**}p < 0.01$, $^{***}p < 0.001$, two-tailed, bootstrap test (for the details of results, see **Supplementary Table 4**). For the full region names, please see **Supplementary Fig. 6**.

**Methods:** To detect the state boundaries of concept representation for each run, we used a version of the Hidden Markov Model (HMM) implemented by Baldassano et al.[8]. This HMM version is optimized for the detection of boundaries based on multi-voxel pattern information of brain regions. In the current analysis, we applied the HMM to each participant's single-trial data. As shown in **a**, we scanned a spherical searchlight with a radius of 5 voxels (= 10 mm) across the whole brain with a step size of 4 voxels, resulting in 3,297 spherical searchlights in total. For each searchlight, we detected the segment boundaries using the HMM with different numbers of segments ranging from $k = 2$ to 12. We used $k = 12$ as the upper limit because $k$ larger than 12 produced too many segments comprised of only one trial, and with $k \leq 12$, we were able to keep the numbers of one-trial segment less than 1% of the number of the total segments. We then calculated the average difference in pattern similarity (i.e., spatial correlation) between intra-versus inter-segment trials for each searchlight, each segmentation solution, and for each $k$, and used it as an indicator of how well the segmentation solutions captured the neural state transitions. We selected the $k$ that produced the largest difference in pattern similarity as the optimal $k$ for a certain searchlight, run, and participant. We then added the optimal $k$s from four runs for each participant, filled the value in the $4 \times 4 \times 4$ voxel cube for each searchlight, z-scored the sum of $k$ values across the whole-brain, and applied smoothing with a 3-mm FWHM Gaussian kernel. As the final step, we conducted one-sample $t$-test on the normalized $k$ maps, and thresholded the results with FDR $q < .05$. We also conducted the ROI analysis with bootstrap tests (10,000 samples) whether the normalized optimal $k$ was different from zero.

**Main findings:** Multiple brain regions within the limbic system, including the ventromedial prefrontal cortex (vmPFC), orbitofrontal cortex, medial temporal lobe, and temporal pole (TP) consistently showed more frequent state transitions (i.e., finer-grained semantic segmentation structure) than other brain regions. Bootstrap tests on the 26 regions-of-interest (ROIs) selected from the previous analyses (see **Supplementary Fig. 6**) provided a similar result—the vmPFC and TP showed the largest numbers of segments among all ROIs (**Fig. 4d** and **Supplementary Table 4**), and the hippocampus and amygdala also showed a larger number of segments than other regions. In addition to the limbic areas, brain regions within the visual and ventral attention networks, including the early visual area, lateral occipital cortex (LOC), anterior insula, and mid-cingulate cortex, also showed larger numbers of segments compared to other brain regions. On the other end of the scale, multiple brain regions within the somatomotor, dorsal attention, and default mode networks showed small numbers of segments, including sensorimotor cortex, dorsolateral prefrontal cortex (dlPFC), posterior cingulate cortex (PCC), and temporal parietal junction (TPJ).

When we compared these findings with the results from Baldassano et al.[8] that used exogenous movie stimuli, there were consistent as well as inconsistent patterns of results. The consistent results between Baldassano et al. and the current study include more segments in the visual cortices and less segments in the PCC and the TPJ (which was near the angular gyrus), which can be interpreted as integrating semantic information along the cortical hierarchy, from the unimodal sensory to higher-order trans-modal brain regions[9]. However, different from Baldassano et al., we found that some high-order trans-modal brain regions within the limbic system showed more segments than other regions, including the vmPFC and TP.

Then, we examined whether these larger numbers of segments (i.e., more frequent state transitions) in the limbic cortical and subcortical regions (the vmPFC, TP, hippocampus/amygdala) and in the visual cortices were due to recurrent activations of similar semantic representations over time using the ratio of intra- to inter-segment pattern similarity (**Supplementary Fig. 9**). The results suggest that the large number of segments within the visual cortical regions could be due to the recurrent activations of similar representations, whereas this was not the case for the limbic regions, which showed a low intra-to-inter segment pattern similarity ratio. Lastly, there is also a possibility that these results are simply due to the low signal-to-noise ratio of these limbic regions, and our supplementary analysis results (**Supplementary Fig. 10**) suggest that it might not be the case in our study—the correlation between the ranks of the optimal k and tSNR was not significant (Spearman's $\rho$ = -0.207, $p$ = 0.3092).

Interestingly, the vmPFC and TP were not even included in the analysis in Baldassano et al.[8] because of these regions' low inter-subject synchrony, which has been also reported in other studies[10-12]. Together these findings may suggest these regions' fundamental roles in endogenous cognitive and affective processes, such as storing and retrieving autobiographical memories of personal experiences and spontaneous thought[13,14]. Therefore, these regions are likely to serve as a major source of the idiosyncrasy across individuals, which will be crucial for advancing personalized neuroscience and personalized treatment for psychiatric disorders.

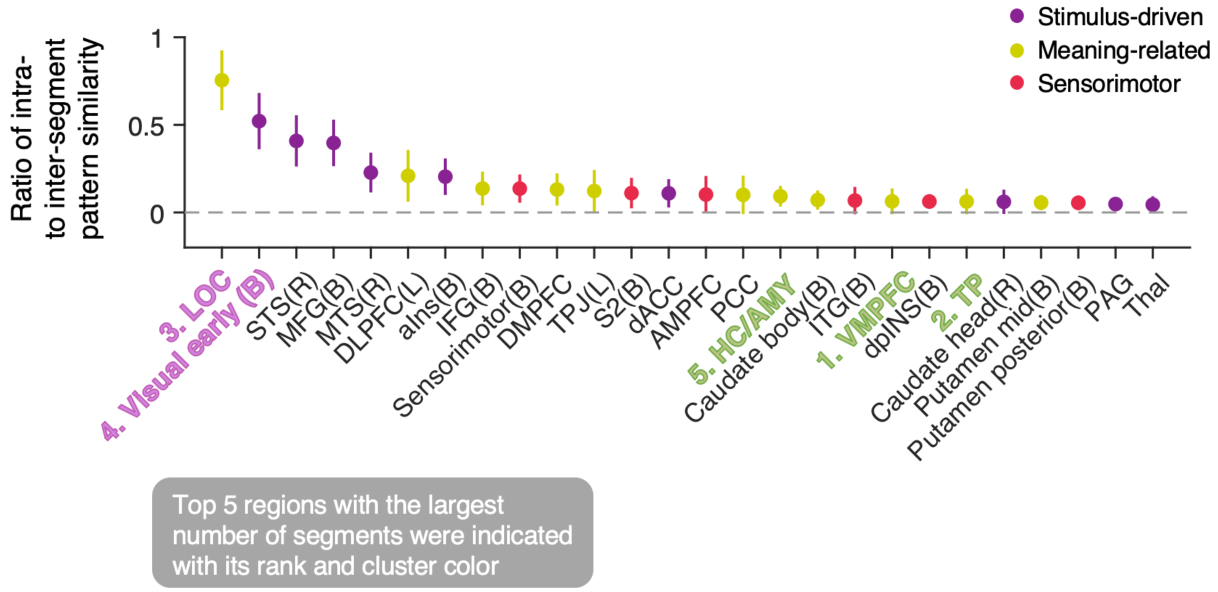


**Supplementary Figure 9. Intra- versus inter-segment pattern similarity of 26 ROIs.** To examine whether the large number of segments (i.e., more frequent state transitions) in some brain regions was due to the recurrent activations of similar semantic representations, we calculated the ratio of intra- to inter-segment pattern similarity. The plot shows the average ratio of intra- versus inter-segment pattern similarity across participants for the 26 ROIs. The error bars represent the standard error of the mean (s.e.m.).

**Main findings:** Among the top 5 regions with the largest number of segments, the LOC and the early visual cortex showed a high intra-to-inter segment pattern similarity ratio, whereas the vmPFC, TP, and the hippocampus/amygdala showed a low intra-to-inter segment pattern similarity ratio. This suggests that the large number of segments within the visual cortical areas could be due to the recurrent activations of similar representations, but it was not the case of the limbic regions.

**Methods:** Based on the segmentation results with the optimal *k* of each region, we calculate the intra-segment pattern similarity using the trials within each segment. We also calculated the inter-segment pattern similarity only considering the trials within the adjacent segments. Then, we divided the intra-state pattern similarity by the inter-state pattern similarity across ROIs and runs and averaged them across 4 runs for each participant.

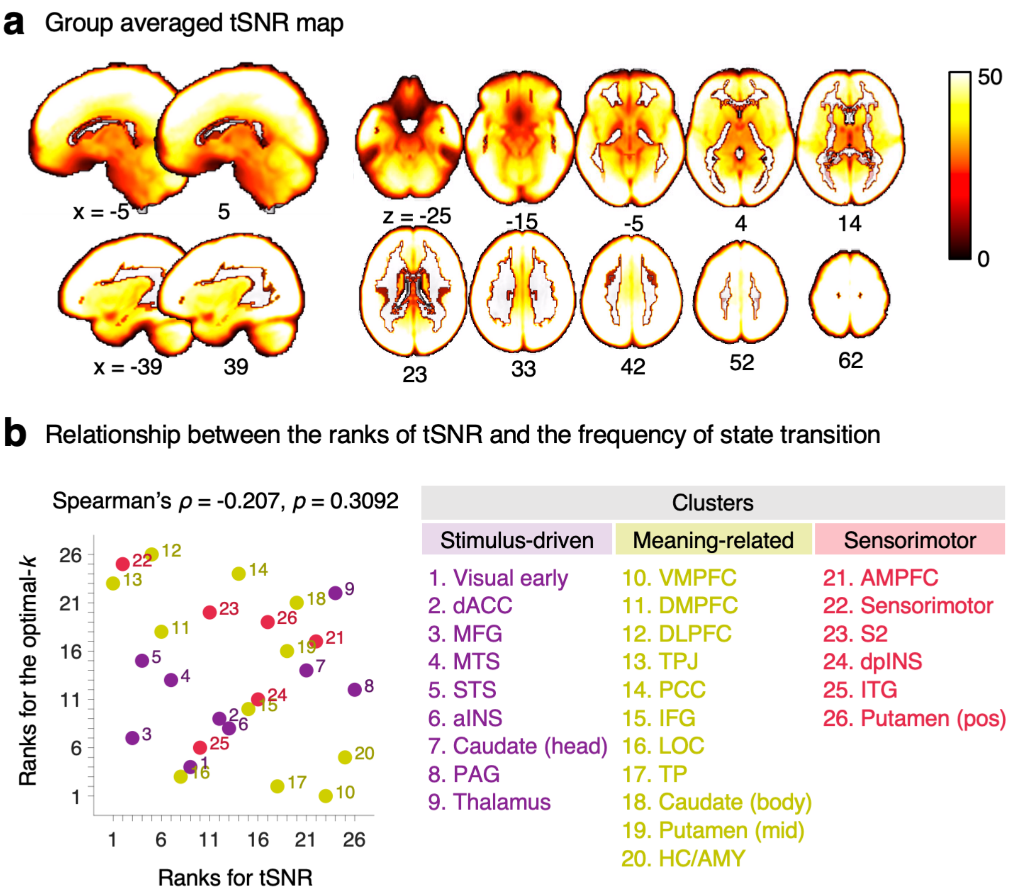


**Supplementary Figure 10. The relationship between the temporal Signal-to-Noise Ratio (tSNR) and the relative frequency of state transition.** We examined whether the results of the relative frequency of state transition were confounded with the levels of signal-to-noise ratio of the BOLD signal. **a**, We calculated the temporal signal-to-noise ratio (tSNR) using the TR images of the concept reflection runs and then averaged the tSNR values across runs and participants ($n$ = 61). The map shows the group-average of the tSNR. **b**, We calculated the Spearman's correlation between the ranks of two variables—the optimal number ($k$) of segments and the tSNR. The correlation between the two ranks was not significant (Spearman's $\rho$ = -0.207, $p$ = 0.3092). For the full region names, please see **Supplementary Fig. 6**.

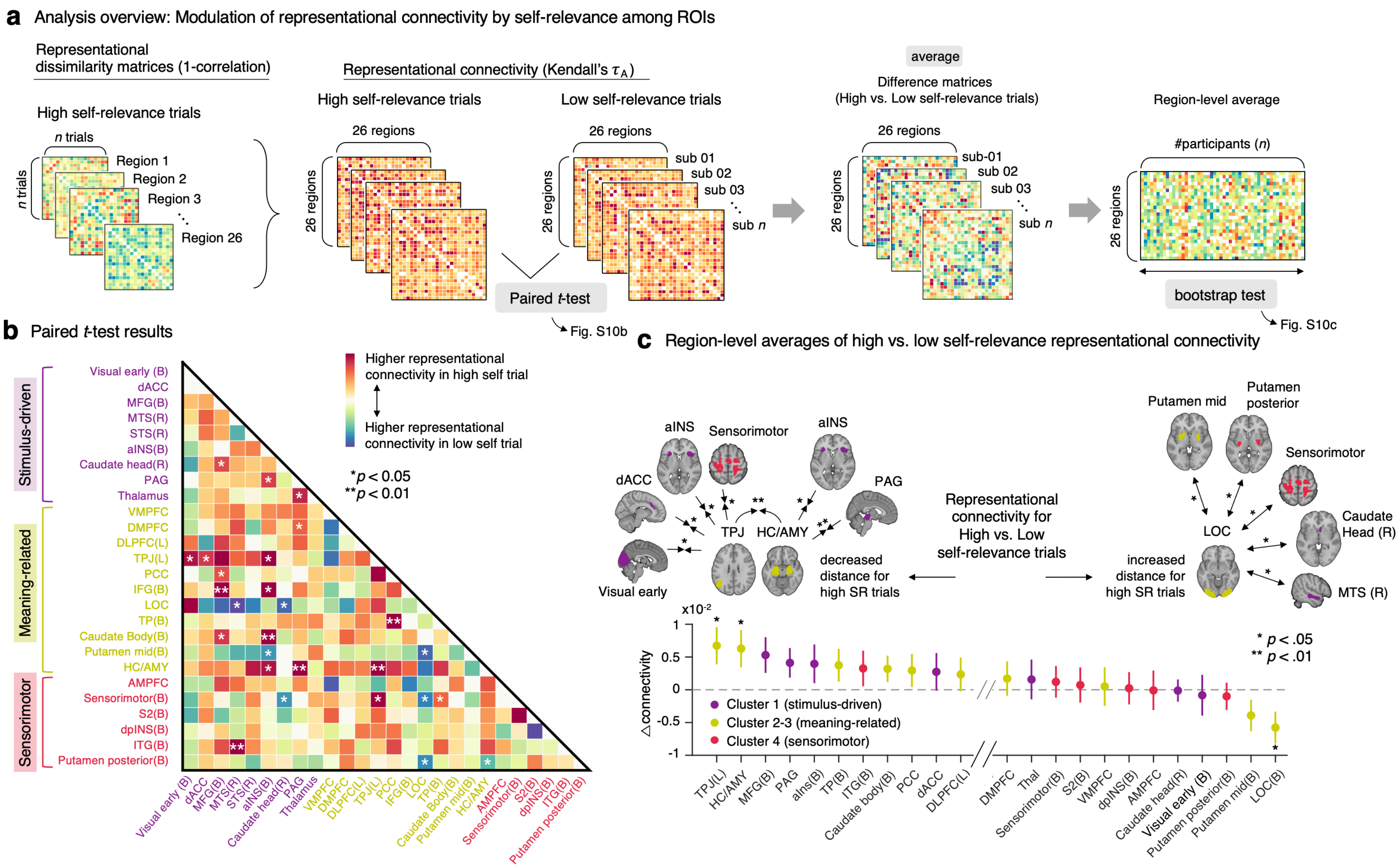


**Supplementary Figure 11. Representational connectivity analysis for the modulatory effects of self-relevance on representational similarity among ROIs. a**, Analysis overview. To identify which brain regions showed the representational changes modulated by the level of self-relevance, we first divided the trials into high versus low self-relevance groups. Then we calculated representational dissimilarity matrices (RDMs) among high self-relevance or low self-relevance trials for each ROI using one minus correlations. With these RDMs, we calculated the representational connectivity among ROIs using Kendall's $\tau_A$[15], resulting in two representational connectivity matrices per participant—one for the high self-relevance condition and the other for the low self-relevance condition. Given that we used 26 ROIs, the size of each representational connectivity matrix was 26 × 26. Using these representational connectivity matrices, we conducted the paired *t*-tests between the high versus low self-relevance conditions. With the difference matrices, we calculated the region-level averages and conducted bootstrap tests (with 10,000 iterations) to identify the ROIs that showed the significant changes in the representational connectivity with other regions by the level of self-relevance.

**b,** The matrix shows the paired *t*-test results with the group average of the difference representational connectivity matrices. The ROI pairs with warm (vs. cold) colors indicate that they showed higher (vs. lower) levels of representational connectivity during high self-relevance than low self-relevance trials. *$p < 0.05$, **$p < 0.01$, uncorrected, two-tailed, bootstrap tests.

**c**, The bottom plot shows the region-level averages of representational connectivity for the high vs. low self-relevance conditions. A dot represents each region, and the y-axis represents the mean difference in representational connectivity for the high versus low self-relevance comparisons. The error bars represent the standard error of the mean (s.e.m.) across individuals. The asterisk indicates the result of bootstrap tests for the paired comparisons, two-tailed.

**Main findings:** The left TPJ, HC/AMY, and bilateral LOC showed significant changes in the representational connectivity with other regions modulated by the level of self-relevance. The left TPJ and HC/AMY showed the overall increases in the representational connectivity with other regions, for the left TPJ, $z = 2.385$, $p = 0.0171$, for the HC/AMY, $z = 2.178$, $p = 0.0294$, whereas the bilateral LOC showed the decreased representational connectivity with other regions, $z = -2.446$, $p = 0.0144$ (for the results of all the ROIs, see **Supplementary Table. 4**). Note that all these regions were a part of the meaning-related cluster.

We then identified the brain regions that showed significant modulations in representational connectivity with these three ROIs. The TPJ and HC/AMY showed decreased representational connectivity with the sensorimotor and salience network brain regions for highly self-relevant trials, including the S1/M1, visual cortex, dACC, and aINS. For the bilateral LOC region, we observed the increased representational connectivity with some basal ganglia regions including putamen and caudate head, S1/M1, and middle temporal sulcus during highly self-relevant trials. Overall, these results suggest that the left TPJ and HC/AMY regions played a role as the hub attractor regions for the high self-relevance trials.

**Supplementary Table 1. Stability of the features across different sets of seed words and test time points (7-week interval)**

| Dynamic features | Valence | | Safety-threat | | Time | | Self-relevance | | Vividness | |
|---|---|---|---|---|---|---|---|---|---|---|
| | *r* | *p* | *r* | *p* | *r* | *p* | *r* | *p* | *r* | *p* |
| Mean | 0.543** | 0.0019 | 0.772*** | 0.0000 | 0.626*** | 0.0002 | 0.609*** | 0.0004 | 0.677*** | 0.0000 |
| Variance | 0.721*** | 0.0000 | 0.754*** | 0.0000 | 0.367* | 0.0462 | 0.693*** | 0.0000 | 0.621*** | 0.0002 |
| <u>*Transition prob.*</u> | | | | | | | | | | |
| Lv.1 → Lv.1 | 0.408* | 0.0254 | 0.271 | 0.1467 | 0.447* | 0.0133 | 0.671*** | 0.0000 | 0.596*** | 0.0005 |
| Lv.2 → Lv.1 | 0.456* | 0.0113 | 0.741*** | 0.0000 | 0.807*** | 0.0000 | 0.440* | 0.0149 | 0.522** | 0.0031 |
| Lv.3 → Lv.1 | 0.589*** | 0.0006 | 0.197 | 0.2967 | -0.075 | 0.6947 | - | - | - | - |
| Lv.1 → Lv.2 | 0.567** | 0.0011 | 0.110 | 0.5621 | 0.314 | 0.0915 | 0.671*** | 0.0000 | 0.596*** | 0.0005 |
| Lv.2 → Lv.2 | 0.579*** | 0.0008 | 0.815*** | 0.0000 | 0.810*** | 0.0000 | 0.440* | 0.0149 | 0.522** | 0.0031 |
| Lv.3 → Lv.2 | 0.489** | 0.0061 | 0.498** | 0.0051 | -0.263 | 0.1607 | - | - | - | - |
| Lv.1 → Lv.3 | 0.505** | 0.0044 | 0.164 | 0.3871 | 0.466** | 0.0095 | - | - | - | - |
| Lv.2 → Lv.3 | 0.438* | 0.0154 | 0.747*** | 0.0000 | 0.755*** | 0.0000 | - | - | - | - |
| Lv.3 → Lv.3 | 0.554** | 0.0015 | 0.661*** | 0.0001 | 0.023 | 0.9020 | - | - | - | - |
| <u>*Steady state prob.*</u> | | | | | | | | | | |
| Lv.1 | 0.690*** | 0.0000 | 0.682*** | 0.0000 | 0.812*** | 0.0000 | 0.565** | 0.0011 | 0.591*** | 0.0006 |
| Lv. 2 | 0.673*** | 0.0000 | 0.757*** | 0.0000 | 0.788*** | 0.0000 | 0.565** | 0.0011 | 0.591*** | 0.0006 |
| Lv. 3 | 0.566** | 0.0011 | 0.791*** | 0.0000 | 0.467** | 0.0092 | - | - | - | - |

*Note.* We examined the stability and test-retest reliability of the Markov chain-based dynamic features across different sets of seed words and across two different time points (7-week interval on average) with a subset of participants ($n$ = 30). The details of how we defined the states and calculated the transition and steady state probabilities, please refer to Methods. Note that since the self-relevance and vividness dimensions had only two discrete states, their correlation values had the same values (e.g., for Lv.1 → Lv.1 and Lv.1 → Lv.2; because one probability is one minus the other probability). One-sample $t$-test for correlations was performed. Lv.1 represents negative, threat, past, low, and low for valence, safety-threat, time, self-relevance, and vividness, respectively. Lv.2 represents neutral,

neutral, present, high, and high for valence, safety-threat, time, self-relevance, and vividness, respectively. Lv. 3 represents positive, safety, and future for valence, safety-threat, and time, respectively. $^{*}p < .05$, $^{**}p < .01$, $^{***}p < .001$, *t*-test for Pearson's correlation, two-tailed.

**Supplementary Table 2. Test-retest reliability and factor loadings of self-report questionnaires**

| Questionnaires | Subscales | FAST-fMRI ($n$ = 30) Test-retest reliability | | FAST-fMRI ($n$ = 62) | | FAST-web ($n$ = 117) | |
|---|---|---|---|---|---|---|---|
| | | | | Factor 1 | Factor 2 | Factor 1 | Factor 2 |
| | | $r$ | $p$ | Negative affect | Positive affect | Negative affect | Positive affect |
| PANAS | Positive affect | 0.278 | 0.1371 | 0.081 | **0.939** | 0.261 | **0.751** |
| PANAS | Negative affect | 0.545 ** | 0.0018 | **0.736** | 0.264 | **1.127** | 0.496 |
| CES-D | Total score | 0.655 *** | 0.0001 | **0.667** | -0.329 | **0.775** | -0.108 |
| MASQ30 | General distress | 0.714 *** | 0.0000 | **0.870** | -0.073 | - | - |
| MASQ30 | Anhedonic depression[a] | 0.415 * | 0.0227 | -0.094 | **0.809** | - | - |
| MASQ30 | Anxiety arousal | 0.723 *** | 0.0000 | **0.571** | 0.323 | - | - |
| STAI-T | Total score | 0.850 *** | 0.0000 | **0.794** | -0.249 | **0.663** | -0.304 |
| RRS | Brooding | 0.563 ** | 0.0012 | **0.747** | 0.060 | - | - |
| RRS | Depressive rumination | 0.562 ** | 0.0012 | **0.814** | 0.070 | **0.547** | -0.110 |
| SIQ | Sum of two items | - | - | - | - | **0.144** | -0.403 |
| LS | Total score | - | - | - | - | **0.306** | -0.511 |
| PWB | Environmental mastery | - | - | - | - | -0.192 | **0.436** |
| PWB | Autonomy | - | - | - | - | 0.081 | **0.486** |
| PWB | Positive relations | - | - | - | - | -0.206 | **0.574** |
| PWB | Purpose in life | - | - | - | - | 0.173 | **0.605** |
| PWB | Personal growth | - | - | - | - | 0.102 | **0.649** |
| PWB | Self-acceptance | - | - | - | - | -0.192 | **0.675** |
| SWLS | Total score | - | - | - | - | -0.138 | **0.545** |

*Note.* Through the FAST-fMRI study, we examined the test-retest reliability of the self-report questionnaires with a 7-week interval using Pearson's correlations. All the questionnaires except for the PANAS-positive affect subscale showed medium to high levels of test-retest reliability, suggesting that these questionnaires provide trait measures. To obtain general negative affectivity score to use it as an outcome variable in predictive modeling, we conducted factor analyses for a two-factor model. The factor analyses were done separately for the FAST-fMRI and FAST-web studies because these two studies conducted different sets of self-report questionnaires—e.g., we included more questionnaires related to positive affectivity in the FAST-web study. The values in bold indicates the higher factor loadings between two factors to show which factor the questionnaire belongs to. PANAS, Positive and Negative Affect Schedule; CES-D, Center for Epidemiologic Studies Depression; MASQ30, 30-item Mood and Anxiety Symptom Questionnaire; STAI-T, State-trait anxiety inventory-Trait version; RRS, Rumination Response Scale; SIQ, Suicidal Ideation Questionnaire; LS, Loneliness Scale; PWB, Psychological Well-Being Scale; SWLS, Satisfaction With Life Scale. $^{*}p < .05$, $^{**}p < .01$, $^{***}p < .001$, $t$-test for Pearson's correlation, two-tailed. [a] reverse coding.

**Supplementary Table 3. Correlation between general negative affectivity and Markov-chain dynamic features**

| Dynamic features | Valence | | Safety-threat | | Time | | Self-relevance | | Vividness | |
|---|---|---|---|---|---|---|---|---|---|---|
| | *r* | *p* | *r* | *p* | *r* | *p* | *r* | *p* | *r* | *p* |
| Mean | -0.432*** | 0.0004 | -0.265* | 0.0374 | 0.137 | 0.2880 | 0.348** | 0.0056 | 0.270* | 0.0338 |
| Variance | 0.343** | 0.0063 | 0.337** | 0.0074 | 0.159 | 0.2175 | 0.005 | 0.9702 | 0.140 | 0.2773 |
| <u>*Transition prob.*</u> | | | | | | | | | | |
| Lv.1 → Lv.1 | 0.294* | 0.0203 | 0.202 | 0.1149 | 0.005 | 0.9682 | -0.213 | 0.0962 | -0.133 | 0.3042 |
| Lv.2 → Lv.1 | 0.360** | 0.0041 | 0.399** | 0.0013 | 0.004 | 0.9767 | -0.258* | 0.0425 | -0.133 | 0.3030 |
| Lv.3 → Lv.1 | 0.416*** | 0.0008 | 0.344** | 0.0062 | -0.124 | 0.3381 | | | | |
| Lv.1 → Lv.2 | -0.173 | 0.1796 | -0.119 | 0.3561 | -0.009 | 0.9432 | 0.213 | 0.0962 | 0.133 | 0.3042 |
| Lv.2 → Lv.2 | -0.045 | 0.7302 | -0.200 | 0.1191 | -0.057 | 0.6603 | 0.258* | 0.0425 | 0.133 | 0.3030 |
| Lv.3 → Lv.2 | -0.207 | 0.1060 | -0.156 | 0.2268 | -0.005 | 0.9697 | | | | |
| Lv.1 → Lv.3 | -0.221 | 0.0843 | -0.109 | 0.4008 | 0.011 | 0.9312 | | | | |
| Lv.2 → Lv.3 | -0.229 | 0.0739 | -0.030 | 0.8189 | 0.114 | 0.3792 | | | | |
| Lv.3 → Lv.3 | -0.180 | 0.1607 | -0.068 | 0.5975 | 0.137 | 0.2892 | | | | |
| <u>*Steday state prob.*</u> | | | | | | | | | | |
| Lv. 1 | 0.484*** | 0.0001 | 0.415*** | 0.0008 | -0.027 | 0.8332 | -0.278* | 0.0288 | -0.186 | 0.1486 |
| Lv. 2 | -0.193 | 0.1327 | -0.249 | 0.0511 | -0.028 | 0.8293 | 0.278* | 0.0288 | 0.186 | 0.1486 |
| Lv. 3 | -0.232 | 0.0702 | -0.001 | 0.9932 | 0.107 | 0.4058 | | | | |

*Note.* Correlation values between the general negative affectivity scores from the factor analysis and the dynamic features from the Markov chain analysis ($n$ = 62). Note that since the self-relevance and vividness dimensions had only two discrete states, some correlation values are same (e.g., for Lv.1 → Lv.1 and Lv.1 → Lv.2; because one probability is one minus the other probability). One-sample $t$-test for correlations was performed. Lv.1 represents negative, threat, past, low, and low for valence, safety-threat, time, self-relevance, and vividness, respectively. Lv.2 represents neutral, neutral, present, high, and high for valence, safety-threat, time, self-relevance, and vividness, respectively. Lv. 3 represents positive, safety, and future for valence, safety-threat, and time, respectively. $^{*}p < .05$, $^{**}p < .01$, $^{***}p < .001$, *t*-test for Pearson's correlation, two-tailed.

**Supplementary Table 4. Optimal number of state segments and representational connectivity for the 26 regions-of-interest (ROIs)**

| ROIs | Relative frequency of state transition | | | Representational connectivity (Kendall's $\tau_A$) | | | |
|---|---|---|---|---|---|---|---|
| | | | | region-level averages | | bootstrap test results | |
| | Average z-score | *z* (boot) | *p* (boot) | high self-relevance | low self-relevance | *z* | *p* |
| *Stimulus-driven cluster* | | | | | | | |
| Visual early (B) | 0.065 | 2.808 ** | 0.0050 | 0.078 | 0.079 | -0.298 | 0.7657 |
| dACC | 0.044 | 0.719 | 0.4719 | 0.068 | 0.065 | 0.956 | 0.3391 |
| MFG(B) | 0.051 | 1.568 | 0.1168 | 0.107 | 0.102 | 1.959 | 0.0501 |
| MTS(R) | 0.028 | 0.576 | 0.5643 | 0.092 | 0.090 | 0.712 | 0.4763 |
| STS(R) | 0.007 | 0.171 | 0.8640 | 0.101 | 0.100 | 0.631 | 0.5279 |
| aINS(B) | 0.044 | 1.251 | 0.2109 | 0.111 | 0.107 | 1.355 | 0.1755 |
| Caudate head(R) | 0.017 | 0.229 | 0.8188 | 0.043 | 0.043 | -0.070 | 0.9441 |
| PAG | 0.028 | 0.636 | 0.5250 | 0.058 | 0.054 | 1.839 | 0.0660 |
| Thalamus | -0.069 | -1.762 | 0.0780 | 0.058 | 0.057 | 0.501 | 0.6164 |
| *Meaning-related cluster* | | | | | | | |
| VMPFC | 0.166 | 4.239 *** | 0.0000 | 0.101 | 0.101 | 0.161 | 0.8717 |
| DMPFC | -0.015 | -0.452 | 0.6514 | 0.131 | 0.129 | 0.628 | 0.5298 |
| DLPFC(L) | -0.179 | -4.795 *** | 0.0000 | 0.104 | 0.102 | 0.918 | 0.3586 |
| TPJ(L) | -0.088 | -1.965 * | 0.0495 | 0.121 | 0.115 | 2.385 * | 0.0171 |
| PCC | -0.128 | -2.221 * | 0.0264 | 0.085 | 0.082 | 1.168 | 0.2426 |
| IFG(B) | 0.042 | 1.334 | 0.1821 | 0.120 | 0.118 | 0.781 | 0.4349 |
| LOC | 0.140 | 3.906 ** | 0.0001 | 0.051 | 0.057 | -2.446 * | 0.0144 |
| TP(B) | 0.148 | 2.846 ** | 0.0044 | 0.077 | 0.073 | 1.510 | 0.1310 |
| Caudate Body(B) | -0.048 | -1.238 | 0.2159 | 0.065 | 0.062 | 1.589 | 0.1121 |
| Putamen mid(B) | -0.006 | -0.226 | 0.8215 | 0.071 | 0.075 | -1.668 | 0.0953 |
| HC/AMY | 0.064 | 2.465 * | 0.0137 | 0.101 | 0.095 | 2.178 * | 0.0294 |
| *Sensorimotor cluster* | | | | | | | |
| AMPFC | -0.007 | -0.101 | 0.9192 | 0.090 | 0.090 | -0.043 | 0.9658 |
| Sensorimotor(B) | -0.133 | -6.389 *** | 0.0000 | 0.100 | 0.099 | 0.501 | 0.6161 |
| S2(B) | -0.047 | -1.379 | 0.1679 | 0.088 | 0.087 | 0.281 | 0.7791 |
| dpINS(B) | 0.030 | 1.225 | 0.2207 | 0.098 | 0.098 | 0.086 | 0.9314 |
| ITG(B) | 0.058 | 1.572 | 0.1159 | 0.096 | 0.093 | 1.212 | 0.2255 |
| Putamen posterior(B) | -0.030 | -0.718 | 0.4725 | 0.063 | 0.064 | -0.508 | 0.6115 |
| Caudate head(R) | 0.017 | 0.229 | 0.8188 | 0.043 | 0.043 | -0.070 | 0.9441 |
| PAG | 0.028 | 0.636 | 0.5250 | 0.058 | 0.054 | 1.839 | 0.0660 |
| Thalamus | -0.069 | -1.762 | 0.0780 | 0.058 | 0.057 | 0.501 | 0.6164 |

*Note.* ROI-level analyses of the Hidden Markov Model (HMM) and representational connectivity. For details of these analyses, please refer to Methods (HMM) and Supplementary

Fig. 10 (representational connectivity analysis). $^{*}p < .05$, $^{**}p < .01$, $^{***}p < .001$, bootstrap tests with 10,000 iteration, two-tailed.

**Supplementary Table 5. Correlations with verbal fluency**

| Variables | r | P | Valence | | Safety-threat | | Time | | Self-relevance | | Vividness | |
|---|---|---|---|---|---|---|---|---|---|---|---|---|
| | | | r | P | r | P | r | P | r | P | r | P |
| Model prediction (predicted negative affectivity) | -0.129 | 0.3169 | | | | | | | | | | |
| # of unique words | 0.299 * | 0.0183 | | | | | | | | | | |
| Mean | | | -0.060 | 0.6439 | -0.151 | 0.2399 | -0.082 | 0.5268 | -0.178 | 0.1652 | -0.144 | 0.2658 |
| Variance | | | -0.081 | 0.5308 | -0.104 | 0.4204 | -0.002 | 0.9895 | 0.176 | 0.1716 | 0.202 | 0.1147 |
| <u>*Transition prob.*</u> | | | | | | | | | | | | |
| Lv.1 → Lv.1 | | | 0.123 | 0.3420 | 0.066 | 0.6108 | 0.053 | 0.6827 | 0.083 | 0.5225 | 0.039 | 0.7649 |
| Lv.2 → Lv.1 | | | -0.077 | 0.5538 | -0.016 | 0.9015 | -0.031 | 0.8109 | 0.306 * | 0.0154 | 0.177 | 0.1684 |
| Lv.3 → Lv.1 | | | -0.038 | 0.7722 | -0.012 | 0.9274 | 0.008 | 0.9495 | | | | |
| Lv.1 → Lv.2 | | | -0.090 | 0.4863 | 0.019 | 0.8823 | 0.023 | 0.8615 | -0.083 | 0.5225 | -0.039 | 0.7649 |
| Lv.2 → Lv.2 | | | 0.058 | 0.6567 | 0.064 | 0.6188 | 0.021 | 0.8717 | -0.306 * | 0.0154 | -0.177 | 0.1684 |
| Lv.3 → Lv.2 | | | 0.197 | 0.1249 | 0.169 | 0.1891 | 0.109 | 0.3986 | | | | |
| Lv.1 → Lv.3 | | | -0.061 | 0.6354 | -0.142 | 0.2714 | -0.233 | 0.0681 | | | | |
| Lv.2 → Lv.3 | | | -0.016 | 0.8991 | -0.084 | 0.5184 | 0.011 | 0.9352 | | | | |
| Lv.3 → Lv.3 | | | -0.141 | 0.2730 | -0.159 | 0.2171 | -0.132 | 0.3075 | | | | |
| <u>*Steady state prob.*</u> | | | | | | | | | | | | |
| Lv. 1 | | | 0.014 | 0.9166 | 0.033 | 0.7964 | 0.001 | 0.9913 | 0.209 | 0.1033 | 0.146 | 0.2565 |
| Lv. 2 | | | 0.086 | 0.5072 | 0.110 | 0.3941 | 0.054 | 0.6767 | -0.209 | 0.1033 | -0.146 | 0.2565 |
| Lv. 3 | | | -0.110 | 0.3950 | -0.167 | 0.1938 | -0.111 | 0.3892 | | | | |

*Note.* We assessed participants' verbal fluency prior to the fMRI experiment to examine whether the individual differences in verbal fluency were related to their FAST performance and results. For the verbal fluency test, we asked participants to produce as many words as possible starting with a Korean alphabet within one minute. We tested with three Korean alphabet letters (ㄱ, ㅇ, and ㅅ). We also asked participants to produce as many animals as possible within one minute. We used the total number of words the participants produced as a verbal fluency score. The verbal fluency score did not show significant correlations with most of the variables except for two variables—the number of unique words in the FAST response and the transition probability of Lv.2 → Lv.1 (or Lv.2) on the self-relevance dimension ($p$ = 0.0183 and 0.0154, respectively; non-significant after the correction for multiple comparisons). $^{*}p < .05$, $t$-test for Pearson's correlation, two-tailed.
